# Biosensor-guided evolution of chalcone synthase enhances biosynthesis of natural and non-natural flavanones

**DOI:** 10.64898/2026.06.12.731874

**Authors:** Erik K. R. Hanko, Chatchai Kesornpun, Joshua N. Whitehead, Reynard Spiess, Christopher J. Robinson, Nigel S. Scrutton

## Abstract

Flavonoids constitute a large class of natural products widely investigated for their bioactive properties, with microbial production offering a potentially scalable alternative to plant extraction. However, achieving structural diversification of these compounds in microbial systems remains challenging, as modification of the flavonoid B-ring typically relies on downstream tailoring enzymes. An alternative strategy is to exploit the intrinsic promiscuity of the canonical flavanone biosynthesis pathway to introduce structural variation at an early stage. Here, we sought to improve microbial production of diverse flavanones by systematically leveraging pathway promiscuity. By constructing a combinatorial library of pathways comprising 4-coumarate-CoA ligase, chalcone synthase, and chalcone isomerase, we enabled the conversion of a panel of ring-substituted cinnamic acid precursors into ten natural and non-natural flavanones. In parallel, we established a genetically encoded biosensor based on the transcriptional regulator FdeR and demonstrated its responsiveness across all ten flavanones. Leveraging this biosensor for high-throughput screening, we performed directed evolution of chalcone synthases from *Hordeum vulgare* and *Arabidopsis thaliana*, identifying enzyme variants that led to improved production of *O*-methylated flavanones, including isosakuranetin, hesperetin, and homoeriodictyol, as well as fluoro-substituted flavanones. In addition, we demonstrated that specific variants of *H. vulgare* chalcone synthase promoted the formation of isoferuloyl-derived derailment products. Collectively, this work establishes the FdeR-based biosensor as a versatile platform for pathway and enzyme engineering, enabling efficient early-stage diversification of flavanones in microbial systems and providing insight into the mutational landscape of chalcone synthases.

## 1. Introduction

Flavonoids constitute a large class of structurally diverse secondary metabolites, with more than 9,000 distinct members reported to date^1^. Predominantly found in plants, they fulfil a wide range of physiological and ecological functions, including roles in UV protection, defence against biotic stress, signalling, reproduction, and development^2, 3^. Owing to the wide spectrum of reported bioactive properties, flavonoids have been extensively investigated for their potential application in nutraceutical, cosmetic, and pharmaceutical fields^2, 4–6^.

Within the flavonoid family, flavanones occupy a central biosynthetic position, acting as branch-point intermediates from which many major flavonoid subclasses arise, including flavones, dihydroflavonols, flavonols, and anthocyanins. In most plants, the canonical flavanone biosynthesis pathway comprises three enzymatic steps: activation of a phenylpropanoid precursor by 4-coumarate-CoA ligase (4CL); formation of a chalcone intermediate by chalcone synthase (CHS), which utilises three units of malonyl-CoA per flavanone molecule; and stereospecific intramolecular cyclisation of the chalcone to yield a chiral flavanone, catalysed by chalcone isomerase (CHI). The resulting flavanone scaffold comprises two aromatic rings (A and B) arranged around a heterocyclic C-ring (Figure 1).

**Figure 1.**
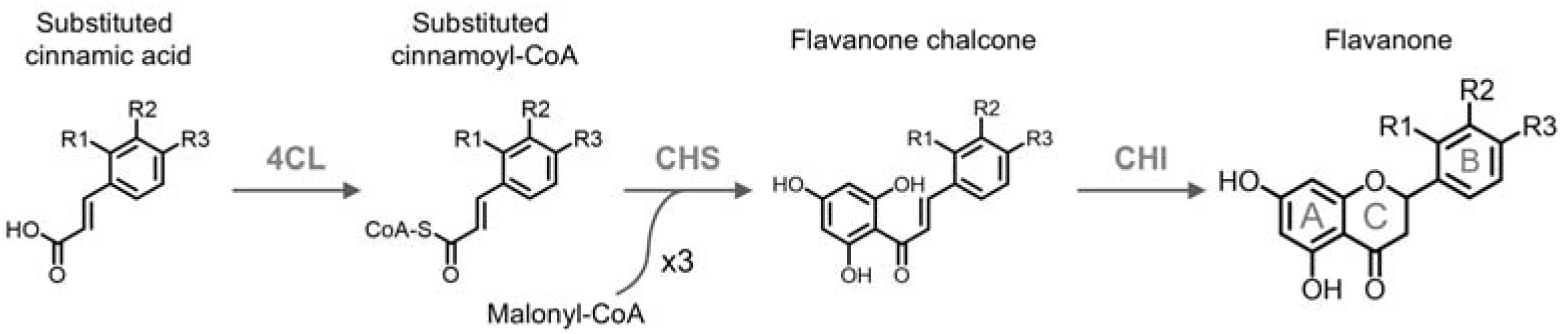
The canonical flavanone biosynthesis pathway. Enzyme abbreviations: 4CL, 4-coumarate-CoA ligase; CHS, chalcone synthase; CHI, chalcone isomerase. The flavanone scaffold consists of two aromatic rings (A and B) arranged around a heterocyclic C-ring. Substituents and the corresponding cinnamic acid and flavanone derivatives are listed in Table 1.

**Table 1.** Flavanones investigated in this study and their corresponding ring-substituted cinnamic acid precursors. Substituent positions (R1–R3) are defined in Figure 1.

| R1 | R2 | R3 | Ring-substituted cinnamic acid substrate<br>systematic or common name | Flavanone product systematic or<br>common name |
| --- | --- | --- | --- | --- |
| H | H | H | Cinnamic acid | Pinocembrin |
| OH | H | H | 2-Hydroxycinnamic acid | 5,7,2'-Trihydroxyflavanone |
| H | OH | H | 3-Hydroxycinnamic acid | 5,7,3'-Trihydroxyflavanone |
| H | H | OH | *p*-Coumaric acid | Naringenin |
| F | H | H | 2-Fluorocinnamic acid | 2'-Fluoro-5,7-dihydroxyflavanone |
| H | F | H | 3-Fluorocinnamic acid | 3'-Fluoro-5,7-dihydroxyflavanone |
| H | H | F | 4-Fluorocinnamic acid | 4'-Fluoro-5,7-dihydroxyflavanone |
| H | H | OCH<sub>3</sub> | 4-Methoxycinnamic acid | Isosakuranetin |
| H | OH | OH | Caffeic acid | Eriodictyol |
| H | OH | OCH<sub>3</sub> | Isoferulic acid | Hesperetin |
| H | OCH<sub>3</sub> | OH | Ferulic acid | Homoeriodictyol |

Microbial biosynthesis of flavonoids offers a potentially scalable alternative to plant extraction, alleviating limitations associated with low and variable product yields in natural sources, while enabling rapid and iterative optimisation of biosynthetic pathways^7–9^. Over the past decade, substantial progress has been made in the microbial production of key flavanones from central metabolism, most notably naringenin and pinocembrin. For example, naringenin titres exceeding 1 g/L have been reported in *Saccharomyces cerevisiae*^10, 11^, while pinocembrin production above 0.5 g/L has been demonstrated in *Escherichia coli*^12^. Access to these gatekeeper flavanones at high titres provides an essential foundation for downstream structural diversification.

Despite these advances, modification of the flavonoid B-ring remains challenging in microbial systems. Incorporation of additional tailoring enzymes, such as cytochrome P450 monooxygenases or flavonoid *O*-methyltransferases for B-ring hydroxylation or *O*-methylation, often introduces new constraints related to heterologous expression, cofactor requirements, or substrate specificity^13–16^. An alternative strategy is to exploit the intrinsic promiscuity of enzymes within the flavanone biosynthesis pathway itself. By utilising ring-substituted cinnamic acids as pathway substrates, this approach enables direct access to B-ring modified flavanones without the need for specialised flavonoid-modifying enzymes^17, 18^. It also provides access to flavanones for which corresponding B-ring tailoring enzymes are not available, including halogenated flavanones, which have been reported to exhibit altered pharmacological properties such as enhanced bioavailability and activity^19^, thereby motivating interest in their biosynthesis.

In this context, chalcone synthase represents a key control point governing entry into flavanone biosynthesis, as it competes with multiple downstream pathways for CoA-activated phenylpropanoid precursors, like *p*-coumaroyl-CoA and feruloyl-CoA^20, 21^. Structural and mechanistic studies have shown that CHS gatekeeps substrate acceptance through steric constraints imposed by a buried active-site cavity and a narrow CoA-binding tunnel. Subtle changes in these features can determine whether a given precursor undergoes full tetraketide extension followed by regiospecific Claisen condensation to yield the chalcone. Alternatively, the precursor may be diverted toward derailment products^22–24^. Together, these structural and mechanistic insights provide a basis for engineering access to non-canonical flavanones through targeted modification of CHS.

Directed evolution provides a powerful framework for probing how changes in enzyme sequence influence catalytic performance and substrate scope. However, for enzymes such as chalcone synthase, whose activity has traditionally been quantified using chromatographic methods, its application has long been constrained by screening throughput. This bottleneck can now be alleviated through the integration of genetically encoded biosensors.

Such biosensors couple intracellular metabolite levels to easily quantifiable reporter outputs, such as fluorescence, thereby enabling high-throughput screening of enzyme variants directly *in vivo*. Both riboswitch- and transcription factor-based biosensors have been successfully applied to enhance naringenin biosynthesis. For example, a riboswitch-based sensor was used to identify *Petunia hybrida* CHS variants with improved catalytic efficiency in *E. coli*^25^, while a biosensor based on the transcriptional regulator TtgR enabled screening of *Sophora japonica* CHS variants with improved catalytic activity in *S. cerevisiae*^11^.

In contrast to TtgR, which in its native context controls expression of a multidrug efflux pump in *Pseudomonas putida*^26^, FdeR is a transcriptional regulator originally implicated in flavonoid catabolism in *Herbaspirillum seropedicae*. FdeR has been shown to mediate transcriptional activation in response to multiple flavonoids, including the flavanone naringenin as well as the flavones chrysin, apigenin, and luteolin^27^.

In this study, we first examine the responsiveness of a biosensor based on the transcriptional regulator FdeR across a panel of structurally diverse flavanones differing in B-ring decoration (Table 1). We then investigate how enzyme identity within the canonical flavanone biosynthesis pathway influences the formation of various B-ring modified flavanones by constructing a combinatorial library of pathways comprising 4-coumarate-CoA ligase, chalcone synthase, and chalcone isomerase, and assess whether biosensor output correlates with analytically quantified flavanone titres. Using the FdeR-based biosensor as a screening tool, we subsequently perform directed evolution of chalcone synthases from *Hordeum vulgare* (barley) and *Arabidopsis thaliana* (mouse-ear cress) to improve production of *O*-methylated- and fluoro-substituted flavanones, respectively. Collectively, this work establishes the FdeR-based biosensor as a versatile platform for flavanone pathway and enzyme screening, and provides enhanced CHS variants as starting points for further enzyme and pathway engineering.

## 2. Materials and methods

### 2.1 Bacterial strains, chemicals, and media

*Escherichia coli* NEB 5-alpha (New England Biolabs) was used for plasmid propagation and cloning. *E. coli* strain SBC016072, a derivative of *E. coli* K-12 MG1655 optimised for flavonoid biosynthesis^28^, was employed for all other experiments described in this study. Ring-substituted cinnamic acid and flavanone analytical standards used in this work are listed in Table S1. Stock solutions of ring-substituted cinnamic acids, used as substrates for flavanone biosynthesis, were freshly prepared in ethanol. For routine laboratory procedures, *E. coli* was cultured in lysogeny broth (LB). For validation of the FdeR-based biosensor and for experiments involving flavanone biosynthesis, *E. coli* was cultured in phosphate-buffered Terrific Broth (Formedium) supplemented with 0.4% (w/v) glycerol (hereafter referred to as production medium). When required, antibiotics were added at the following final concentrations: chloramphenicol (25 µg/mL) and kanamycin (50 µg/mL).

### 2.2 Plasmid and strain construction

Oligonucleotide primers were synthesised by Integrated DNA Technologies (IDT) and are listed in Table S2. Gene parts were designed using PartsGenie^29^, with coding sequences codon-optimised for *E. coli* and ribosome binding site translation initiation rates set to 20,000. Gene parts were synthesised by TWIST Bioscience, and their sequences are provided in Table S3. Plasmid DNA was extracted using the QIAprep Spin Miniprep Kit (Qiagen). DNA fragments for cloning were amplified by PCR using the Q5 High-Fidelity 2X Master Mix (NEB) and gel-purified with the Zymoclean Gel DNA Recovery Kit (Zymo Research). Plasmids were constructed either by restriction enzyme-based cloning or by HiFi DNA Assembly. Restriction enzymes, T4 DNA ligase, and the NEBuilder HiFi DNA Assembly Master Mix were purchased from NEB, and all PCR, digestion, ligation, and assembly reactions were carried out according to the manufacturer’s instructions. A detailed description of the construction of each plasmid is provided in the Supplementary Methods. All plasmids were verified by Sanger sequencing (Eurofins Genomics) or whole-plasmid sequencing (Source BioScience). A complete list of plasmids used and generated in this study is provided in Table S4. Plasmids were transformed into chemically competent *E. coli* cells by heat shock^30^.

### 2.3 Strain cultivation

For routine laboratory procedures, *E. coli* was cultured at 37 D with orbital shaking at 200 rpm. For validation of the FdeR-based biosensor and for small-scale flavanone biosynthesis experiments, single colonies of freshly transformed *E. coli* were used to inoculate 1 mL of production medium supplemented with the relevant antibiotics. Cultures were grown in 96-well deep-well plates (DWP) sealed with breathable seals.

Seed cultures were incubated at 30 D and 80% humidity with orbital shaking at 850 rpm for 18 h. Main cultures were prepared by diluting seed cultures 1:200 into fresh production medium (1 mL) in 96-well DWP and incubated under the same conditions. After 6 h of growth, when cultures had reached an OD_600_ of 1.0–2.0, either a flavanone ligand (for biosensor validation) or a ring-substituted cinnamic acid substrate (for flavanone biosynthesis) was added at the appropriate concentration. Optionally, IPTG was added to a final concentration of 0.1 mM to induce expression of flavanone biosynthesis pathway enzymes. Cultures were then returned to the shaker-incubator, and samples were collected after 24 h.

The large-scale production, purification, and structural verification of the flavanones 5,7,2′-trihydroxyflavanone, 2′-fluoro-5,7-dihydroxyflavanone, 3′-fluoro-5,7-dihydroxyflavanone, and 4′-fluoro-5,7-dihydroxyflavanone are described in the Supplementary Methods.

### 2.4 Sample preparation and quantification of target compounds

Samples for analytical confirmation and quantification of target compounds, by ultra-performance liquid chromatography with diode-array detection (UPLC-DAD), liquid chromatography-tandem mass spectrometry (LC-MS/MS), and LC-quadrupole time-of-flight mass spectrometry (LC-QTof MS) analysis, were prepared by transferring 50 µL of the bacterial culture to a 96-well microtitre plate and adding 150 µL methanol. Samples were subsequently stored at −80 D. On the day of analysis, samples were thawed at room temperature, centrifuged for 10 min at 4,000 rpm, and cell-free supernatants were diluted as required with 40% methanol prior to analysis.

Quantification of hesperetin, isosakuranetin, and homoeriodictyol, was performed by LC-MS/MS analysis. All other target compounds were quantified using UPLC-DAD.

LC-MS/MS analysis was carried out using a Waters ACQUITY UPLC I-Class binary system coupled to a Waters Xevo TQ-XS triple quadrupole mass spectrometer, equipped with a Waters ACQUITY BEH C18 column (50 × 2.1 mm, 1.7 µm). The column temperature was maintained at 45 □. Separation was achieved at a flow rate of 0.5 mL/min using a binary mobile phase consisting of solvent A (H_2_O with 0.1% formic acid) and solvent B (acetonitrile with 0.1% formic acid). The gradient elution programme was as follows: 0–1.0 min, 15% B; 1.0–4.5 min, linear gradient from 15% to 95% B; 4.5–4.9 min, held at 95% B; 4.9–5.0 min, return to 15% B; 5.0–6.0 min, re-equilibration at 15% B. Samples were maintained at 10 D in the autosampler, and the injection volume was 1 µL. MS parameters and multiple reaction monitoring (MRM) transitions are provided in Table S5. Peak areas were integrated using Waters MassLynx software.

UPLC-DAD analysis was performed using an Agilent 1290 Infinity II LC system coupled to a diode-array detector measuring absorbance at 275, 290, and 320 nm. The chromatography method mirrored the LC-MS/MS method described above, except that methanol was used as solvent B instead of acetonitrile, and the injection volume was 5 µL. Peak areas were integrated using Agilent OpenLab software. Metabolite concentrations were determined using calibration curves generated from standards of known concentrations.

### 2.5 LC-QTof MS analysis

Accurate-mass measurement and fragmentation analysis were performed using a Waters ACQUITY UPLC I-Class system coupled to a Waters ion mobility quadrupole time-of-flight mass spectrometer (Vion IMS-QToF). Chromatographic separation was carried out using a Waters ACQUITY BEH C18 column (50 × 2.1 mm, 1.7 µm), with the column temperature maintained at 45 D. Separation was achieved at a flow rate of 0.5 mL/min using a binary mobile phase consisting of solvent A (H_2_O with 0.1% formic acid) and solvent B (methanol). The gradient elution programme was as follows: 0–1.0 min, 15% B; 1.0–7.0 min, linear gradient from 15% to 66% B; 7.0–7.1 min, linear gradient from 66% to 95% B; 7.1–9.0 min, held at 95% B; 9.0–9.1 min, return to 15% B; 9.1–10.0 min, re-equilibration at 15% B. Electrospray ionisation was performed in negative ion mode, with an ion source temperature of 130 D. The mass scan range was 50–1000 *m*/*z*, with a scan time of 1 s. Continuous mass calibration was achieved using LockSpray analysis of a reference solution containing leucine enkephalin (50 ng/mL), acquired every 3 min during the chromatographic run. Data were collected in MS^E^ mode, with the instrument alternating between low-energy collision conditions (6 eV) for precursor ion collection and high collision energies (15–40 eV ramp) for product ion generation throughout the run. Data were processed and analysed using Waters UNIFI software. Putative product ions were predicted using Waters UNIFI software and CFM-ID (version 4.0)^31^.

### 2.6 Fluorescence measurements

To quantify red fluorescent protein (RFP) reporter output, 15 µL of bacterial culture were transferred to a 96-well microtitre plate (black with clear bottom; Greiner Bio-One) containing 135 µL of phosphate-buffered saline (PBS). RFP fluorescence and optical density at 600 nm (OD_600_) were measured using a CLARIOstar Plus plate reader (BMG Labtech). Fluorescence excitation and emission wavelengths were set to 585 nm and 620 nm, respectively, with the gain factor fixed at 1,500. To account for medium autofluorescence and background absorbance, fluorescence and OD_600_ values were corrected by subtracting measurements obtained from cell-free diluted medium. Corrected fluorescence values were subsequently normalised to OD_600_ to account for differences in cell density.

### 2.7 Statistical analysis

All statistical analyses, including unpaired two-tailed *t* tests and three-way ANOVA, were performed using software GraphPad Prism (version 10.2.2). A *p* value < 0.05 was considered statistically significant.

## 3. Results and discussion

### 3.1 FdeR responds to a broad range of flavanones

To establish a robust biosensor platform suitable for screening biosynthetic pathways leading to structurally diverse flavanones and for enabling high-throughput directed evolution of enzymes involved in flavanone biosynthesis, we first assessed whether the transcriptional regulator FdeR is capable of sensing flavanones beyond its previously characterised ligand, naringenin^27^. Such breadth of responsiveness is essential if FdeR-based biosensors are to be applied to the discovery and optimisation of pathways yielding both natural and non-natural flavanones.

To test this, we constructed a biosensor circuit comprising the *fdeR* gene and the native intergenic region between *fdeR* and *fdeA* from *Herbaspirillum seropedicae*, which harbours both the constitutive *fdeR* promoter and the inducible *fdeA* promoter^27^. The *fdeA* promoter was transcriptionally linked to an *rfp* reporter gene, such that flavanone-dependent activation of reporter transcription could be quantified via fluorescence output (Figure 2A). The resulting biosensor plasmid was introduced into *Escherichia coli* strain SBC016072, a derivative of *E. coli* KD12 MG1655 optimised for flavonoid biosynthesis^28^.

**Figure 2.**
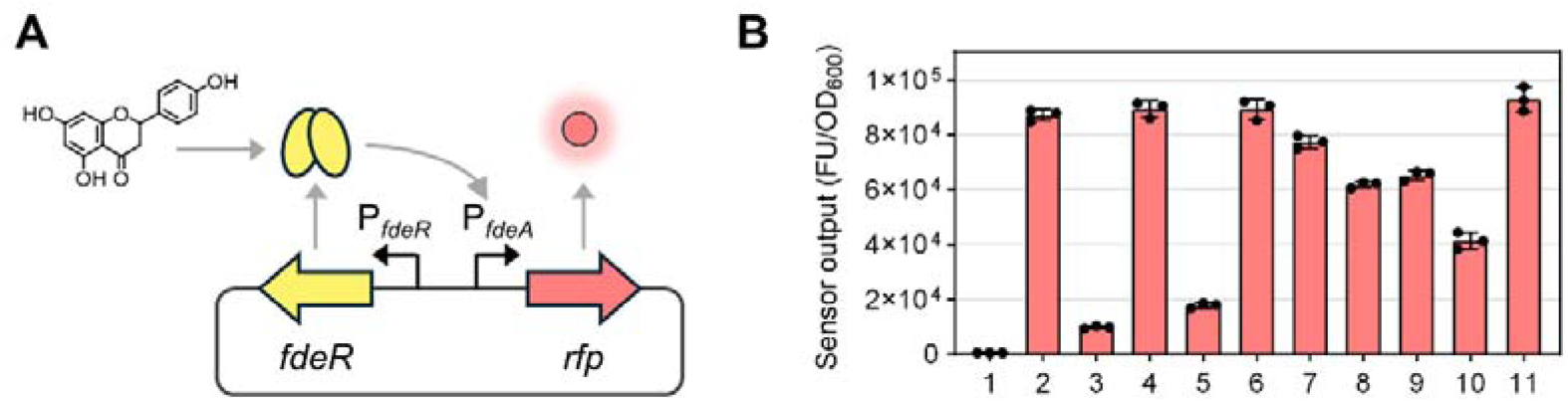
Validation of an FdeRDbased biosensor for diverse flavanones. **A** Schematic representation of the flavanone biosensor circuit in plasmid SBC016239. The transcriptional regulator FdeR is expressed from its native promoter (P*_fdeR_*), while the flavanone-inducible promoter of *fdeA* (P*_fdeA_*) is linked to an *rfp* reporter gene. Binding of a flavanone ligand (e.g. naringenin) to FdeR activates transcription from P*_fdeA_*, resulting in fluorescent reporter output. **B** Fluorescence reporter output of *E. coli* strain SBC016072 harbouring the biosensor plasmid SBC016239, measured 24 h after supplementation with individual flavanones at a final concentration of 0.1 mM. Ligands are shown in the following order: uninduced control (1), pinocembrin (2), 5,7,2′-trihydroxyflavanone (3), naringenin (4), 2′-fluoro-5,7-dihydroxyflavanone (5), 3′-fluoro-5,7-dihydroxyflavanone (6), 4′-fluoro- 5,7-dihydroxyflavanone (7), isosakuranetin (8), eriodictyol (9), hesperetin (10), and homoeriodictyol (11). Data are presented as mean ± standard deviation (*n* = 3).

Biosensor performance was evaluated by supplementing cultures with a panel of ten flavanones at a final concentration of 0.1 mM and quantifying fluorescence output after 24 h. The tested compounds comprised both naturally occurring flavanones, including pinocembrin, 5,7,2′-trihydroxyflavanone, naringenin, isosakuranetin, eriodictyol, hesperetin, and homoeriodictyol, as well as non-natural fluorinated analogues, including 2′-, 3′-, and 4′-fluoro-5,7-dihydroxyflavanone.

All tested flavanones elicited a measurable biosensor response, although the magnitude of reporter induction varied substantially between ligands (Figure 2B). Homoeriodictyol triggered the strongest response, resulting in an approximately 150-fold increase in normalised fluorescence relative to the uninduced control. In contrast, 2′-fluoro-5,7-dihydroxyflavanone produced a more moderate, yet clearly distinguishable, 16-fold induction. The observed variation in reporter output likely reflects differences in ligand–FdeR affinity, ligand uptake, or a combination thereof.

Despite these quantitative differences, the observation that all tested flavanones activated the biosensor demonstrates that FdeR exhibits a notable degree of ligand promiscuity. Importantly, FdeR is able to sense and respond to a range of structurally related flavanones, encompassing both native plant metabolites such as naringenin and non-natural analogues bearing fluorine substitutions. This breadth of responsiveness establishes FdeR as a versatile biosensor suitable for monitoring the biosynthesis of diverse flavanone scaffolds and underpins its application in downstream pathway screening and enzyme engineering efforts.

### 3.2 Optimisation of flavanone biosynthesis across diverse substrates

The microbial production of structurally diverse flavanones has primarily been accomplished via two parallel strategies. One approach focuses on maximising the biosynthesis of the gatekeeper flavanone naringenin, followed by structural modification of the B-ring by dedicated tailoring enzymes^13^. A second approach exploits the inherent promiscuity of the canonical three-enzyme flavanone biosynthesis pathway, utilising ring-substituted cinnamic acids as starter substrates to access less common natural or non-natural flavanones, including fluorinated derivatives^18, 32^. While impressive titres of naringenin have been achieved through enzyme screening and pathway optimisation^10, 11^, its subsequent diversification in microbial hosts remains challenging, often requiring hydroxylation and methylation reactions and yielding complex mixtures of flavonoid products^13^. In contrast, comparatively few studies have leveraged this second strategy, systematically combining pathway optimisation with enzyme promiscuity, to enable efficient biosynthesis of structurally more complex flavanones using only the core 4CL–CHS–CHI pathway^18, 32–34^.

To explore the latter approach, we sought to identify enzyme combinations that enable efficient biosynthesis of a broad range of B-ring-modified flavanones in *E. coli*, spanning non-*O*-methylated, *O*-methylated, and fluorinated compounds. Specifically, we aimed to systematically dissect how enzyme identity at each step of the three-enzyme flavanone biosynthesis pathway influences product formation across a panel of eleven ring-substituted cinnamic acid substrates (Table 1).

To this end, we constructed a combinatorial library of eight flavanone biosynthesis pathways comprising two enzyme homologues at each step of the core pathway (Figure 3A). The library included either 4CL1 (Gm4CL1, UniProt: Q8S564) or 4CL4 (Gm4CL4, UniProt: P31687) from *Glycine max*, either CHS from *Arabidopsis thaliana* (AtCHS, UniProt: P13114) or CHS2 from *Hordeum vulgare* (HvCHS2, UniProt: Q96562), and either CHI2 from *A. thaliana* (AtCHI2, UniProt: Q9FKW3) or CHI1 from *Medicago sativa* (MsCHI1, UniProt: P28012). The gene order and promoter elements were retained from previously optimised pinocembrin- and naringenin-producing constructs^7, 35^. In addition, all pathway plasmids carried the FdeR-based biosensor.

**Figure 3.**
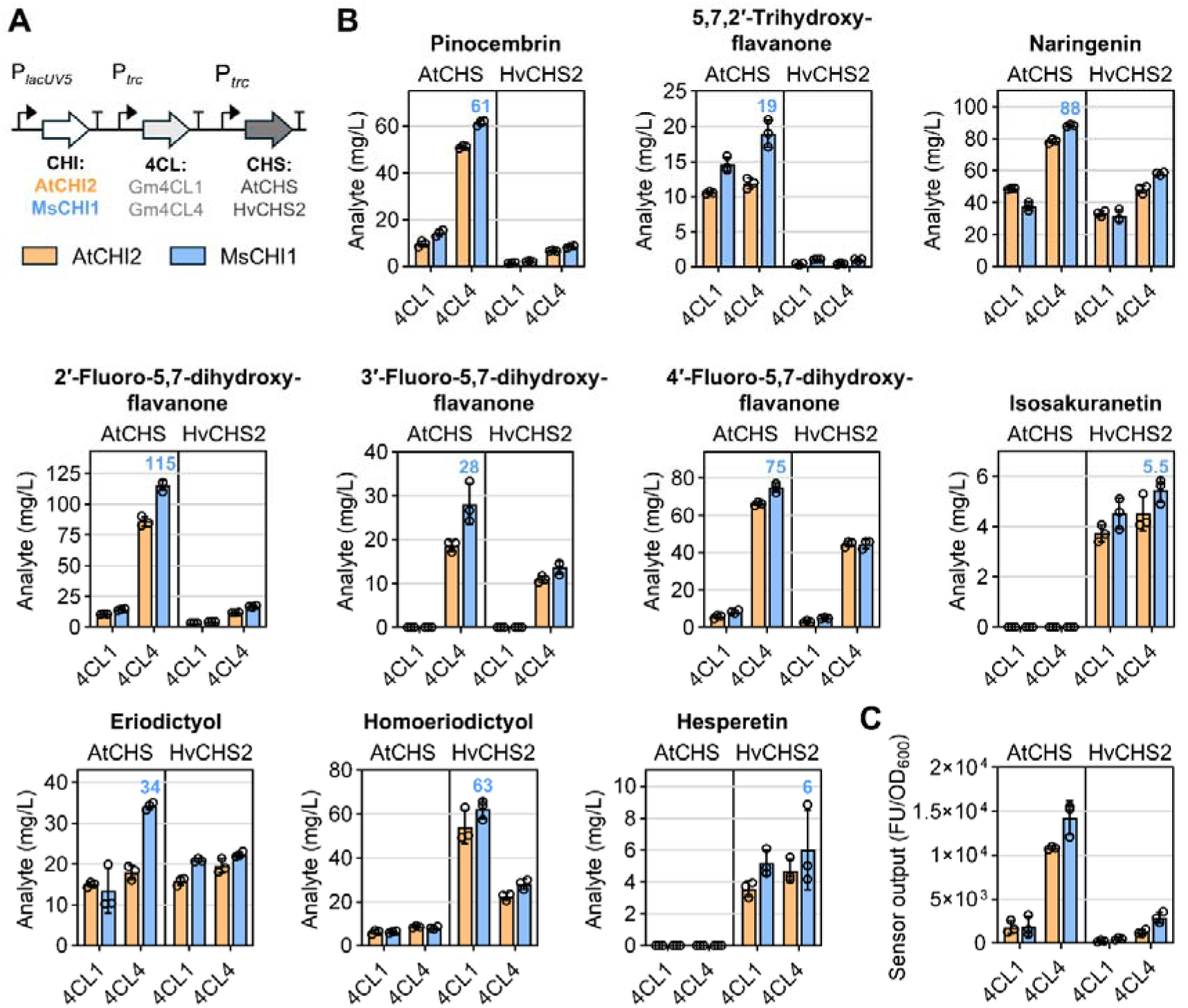
Combinatorial pathway screen for flavanone biosynthesis. **A** Illustration of gene order and promoter elements used to construct the combinatorial pathway library, with each pathway variant comprising one of two homologues for each of 4CL, CHS, and CHI. In addition, all pathway variants carry the FdeR-based biosensor. **B** Flavanone titres obtained in *E. coli* strain SBC016072 carrying the different pathway variants, quantified 24 h after supplementation with the corresponding ring-substituted cinnamic acid at a final concentration of 0.5 mM. The flavanone titre of the highest-producing variant is indicated in each panel. **C** Fluorescence reporter output of *E. coli* strain SBC016072 carrying the same pathway variants, quantified 24 h after supplementation with 3-hydroxycinnamic acid at a final concentration of 0.5 mM. CHS and 4CL identities are indicated within each chart, while CHI identity is represented by bar colour (orange: AtCHI2; blue: MsCHI1). Data are presented as mean ± standard deviation (*n* = 3).

The selection of enzyme homologues was guided by prior observations of substrate scope and pathway performance. The combination of Gm4CL4, AtCHS, and AtCHI2 has been reported to yield high titres of naringenin, pinocembrin, and eriodictyol, but to perform poorly with *O*-methylated substrates such as ferulic acid and 4-methoxycinnamic acid^35^. In contrast, HvCHS2 has been shown to accommodate methoxy-substituted cinnamoyl-CoA intermediates, including feruloyl- and isoferuloyl-CoA^34^. We also included Gm4CL1, which has been reported to accept bulkier ring-substituted cinnamic acid derivatives^36, 37^, and MsCHI1, which has been successfully employed for the production of *O*-methylated flavanones such as homoeriodictyol and hesperetin in related heterologous systems^34^. All eight pathway variants were transformed into *E. coli* strain SBC016072 and evaluated following supplementation with 0.5 mM of the corresponding ring-substituted cinnamic acid. Substrate consumption, product titres (where authentic standards were available), and FdeR-based biosensor output were quantified 24 h after substrate addition.

Across the flavanones for which authentic standards were available, pronounced differences in pathway performance were observed. The pathway comprising Gm4CL4, AtCHS, and MsCHI1 consistently produced the highest titres of all non-*O*-methylated flavanones tested, including pinocembrin, 5,7,2′-trihydroxyflavanone, naringenin, the three fluorinated flavanones, and eriodictyol (Figure 3B). In contrast, production of the *O*-methylated flavanones isosakuranetin and hesperetin was favoured by the combination of Gm4CL4, HvCHS2, and MsCHI1, whereas homoeriodictyol production was highest in the pathway comprising Gm4CL1, HvCHS2, and MsCHI1. Although microbial production of isosakuranetin via *O*-methylation of naringenin has been reported previously^13^, direct formation of isosakuranetin from a methoxy-substituted cinnamic acid starter unit has not, to our knowledge, been described. Furthermore, while fluoro-substituted flavanones have previously been produced at low mg/L titres in yeast^38^, the titres achieved here were considerably higher, reaching up to 115 mg/L and corresponding to a conversion yield of 86% for 2′-fluoro-5,7-dihydroxyflavanone. This highlights the potential of substrate-mediated diversification to access non-natural flavonoid space without the need for additional tailoring enzymes.

To quantify the contribution of individual enzymes to product formation and to identify context-dependent effects, product titres were analysed using a three-way ANOVA with 4CL, CHS, and CHI treated as categorical factors. Rather than comparing individual pathway variants as isolated constructs, this approach enabled systematic decomposition of variance attributable to enzyme identity at each pathway step, as well as to interaction effects between enzymes.

Across the flavanones analysed, the relative contributions of individual enzymes to product formation varied substantially between substrates. For several flavanones, including pinocembrin and 2′-fluoro-5,7-dihydroxyflavanone, CHS identity accounted for a large fraction of the observed variance in product titre, with an additional substantial contribution arising from 4CL–CHS interaction effects (Table S6). These interaction effects reflect context dependence, whereby product formation depends on the specific pairing of starter-unit activation and chalcone-forming activities. In contrast, for naringenin, 3′-, and 4′-fluoro-5,7-dihydroxyflavanone, variance in product titre was dominated by 4CL identity, indicating that activation of the respective ring-substituted cinnamic acid constitutes the primary bottleneck under these conditions. Consistent with this interpretation, pathways containing Gm4CL1 showed neither detectable depletion of 3-fluorocinnamic acid (Figure S1) nor accumulation of the corresponding flavanone product, indicating that this enzyme is unable to efficiently activate this substrate.

A distinct pattern was observed for the *para*-*O*-methylated flavanones isosakuranetin and hesperetin. In these cases, CHS overwhelmingly dominated product formation, accounting for over 85% of the total variance, while contributions from 4CL, CHI, and interaction effects were minimal. These results indicate that, once 4-methoxycinnamic acid or isoferulic acid is successfully activated, flux through the pathway is almost entirely constrained by CHS performance and is relatively insensitive to the choice of 4CL and CHI under the conditions tested. In contrast, eriodictyol did not exhibit a similarly distinct dependence on a single enzymatic step, with most pathway variants yielding comparable product titres. Here, variance was distributed across multiple factors, suggesting that no single enzyme constitutes a dominant bottleneck for eriodictyol biosynthesis within the set of candidate enzymes examined.

Despite frequent statistical significance of CHI-associated effects, CHI identity accounted for only a minor fraction of the total variance in product titre (< 5%) for most flavanones, with the exception of eriodictyol, where CHI contributed a larger, though still not dominant, fraction of the observed variance. This indicates that, within the set of pathway variants examined, CHI primarily plays a modulatory role rather than constituting a major determinant of flux through the flavanone biosynthesis pathway.

Because an authentic standard was not available for 5,7,3′-trihydroxyflavanone, absolute quantification and unambiguous confirmation of its production by chromatographic methods alone was not possible. However, supplementation with 3-hydroxycinnamic acid resulted in a marked increase in FdeR-dependent biosensor output, with up to a 62-fold difference in normalised fluorescence between pathway variants (Figure 3C). This response is consistent with the formation of 5,7,3′-trihydroxyflavanone, or a closely related derivative capable of activating the biosensor.

Finally, to assess whether biosensor output provides a quantitative proxy for flavanone production across the combinatorial pathway variants, we examined the relationship between normalised fluorescence and product titres measured across the eight pathway variants for each substrate. For most flavanones, including pinocembrin, 5,7,2′-trihydroxyflavanone, naringenin, homoeriodictyol, and the fluorinated flavanones, biosensor output correlated strongly with product titre (Figure 4), indicating that FdeR-dependent reporter activation reflects flavanone levels over the range of titres produced. In contrast, weak or no correlation was observed for the three remaining flavanones, most notably hesperetin, for which biosensor output did not scale with measured product titre. This lack of correlation suggests that, although hesperetin is produced by pathway variants containing HvCHS2, intracellular concentrations may remain below the activation threshold of the FdeR-based biosensor. Together, these results demonstrate that while the biosensor provides a robust quantitative readout for many flavanones, its utility is greatest when product titres fall within the sensor’s detection range, which could in principle be extended through adjustment of substrate feeding strategies or through targeted engineering of FdeR to alter its sensitivity toward specific substrates.

**Figure 4.**
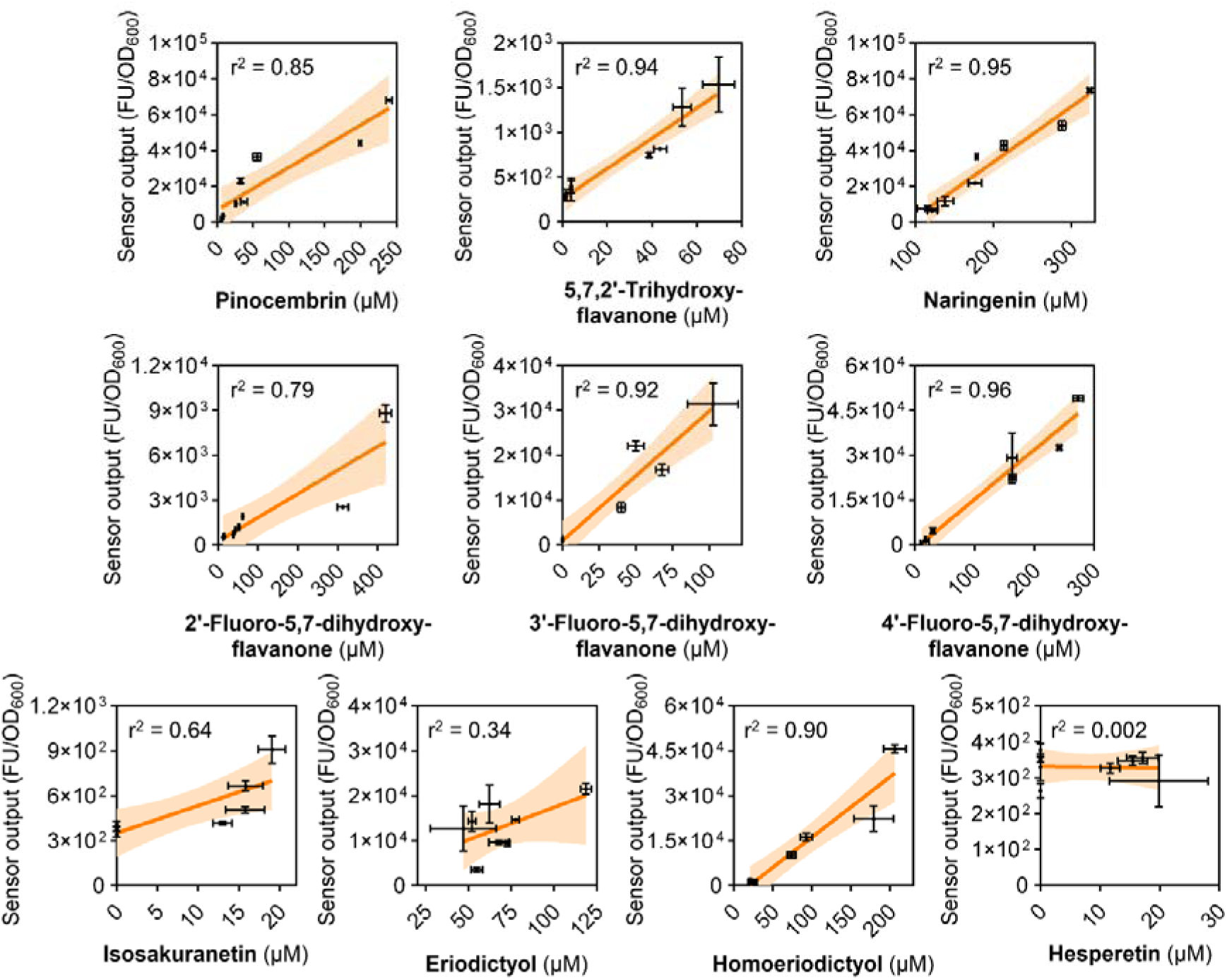
Correlation between biosensor output and flavanone titre within the combinatorial pathway library. Product titres correspond to the data presented in Figure 3. Each data point represents the mean and error bars indicate the standard deviation of three biological replicates. The Pearson correlation coefficient r^2^ is shown (*n* = 8), together with the 95% confidence interval of the linear regression.

### 3.3 Directed evolution of HvCHS2 enhances production of *O*-methylated flavanones

Given the strong dependence of *O*-methylated flavanone biosynthesis on CHS identity observed in the combinatorial pathway analysis, we next targeted HvCHS2 for engineering. While comparatively high titres of the *meta*-*O*-methylated flavanone homoeriodictyol were achieved in the combinatorial pathway screen, production of the *para*-*O*-methylated flavanones isosakuranetin and hesperetin remained low. Previous work has demonstrated that mutation of surface-exposed residues in HvCHS2 (e.g. Q232P and D234V) can enhance production of homoeriodictyol and hesperetin, an effect that was suggested to arise from increased protein abundance and altered substrate affinity^34^. Building on these observations, we sought to systematically explore the mutational landscape of HvCHS2 to improve flavanone biosynthesis from *O*-methylated cinnamoyl-CoA starter units.

Guided by the protein structure of HvCHS2, complexed with CoA and eriodictyol (PDB: 8B3C)^34^, we selected fifteen residues located within the active-site initiation and elongation cavity or along the CoA-binding tunnel for site-saturation mutagenesis (Figure 5A). These residues (T134, S135, M139, Q163, E194, I195, T196, L216, D219, I256, L265, L269, V273, M339, and S340) were chosen based on their proximity to the enzyme-bound starter unit or the growing polyketide chain. We reasoned that substitutions at these positions, particularly to smaller or less polar side chains, could help accommodate more sterically demanding ring-substituted cinnamoyl-CoA derivatives, and reduce unfavourable polar–hydrophobic mismatches introduced by *O*-methylation, which eliminates phenolic hydrogen bond donation and increases local hydrophobicity. In addition, two surface-exposed residues (C204 and D317), which show low sequence conservation among commonly utilised chalcone synthases (Figure S2), were included.

**Figure 5.**
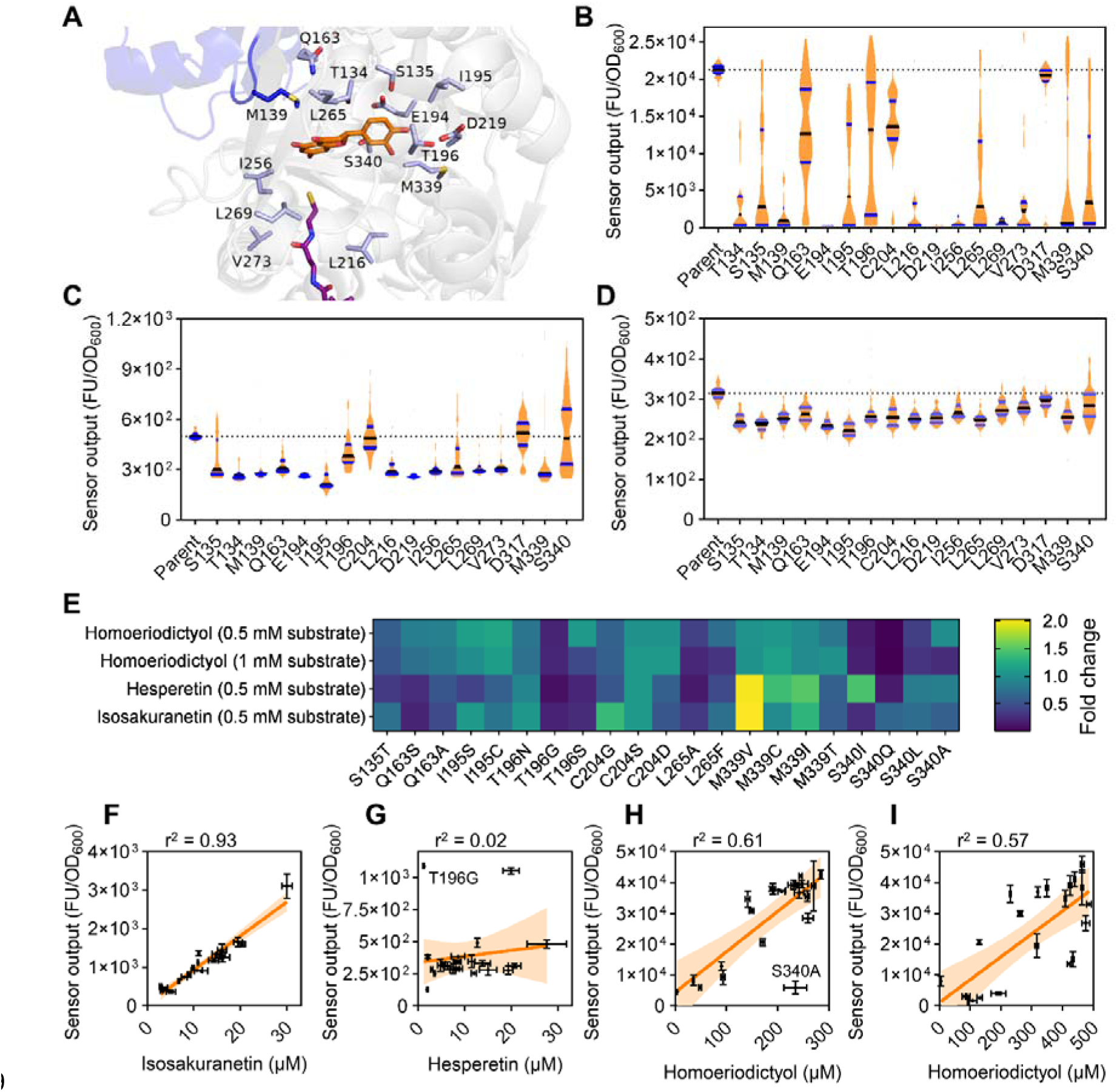
Directed evolution of HvCHS2. **A** Illustration of the active site and substrate channel of HvCHS2 complexed with eriodictyol (orange) and CoA (purple) from PDB: 8B3C. Residues selected for mutation from chain A (grey cartoon) are shown in light blue. M139, which comes from the dimer interface with chain B (blue cartoon) is shown in dark blue. The loop on chain B containing M139 is highlighted. For clarity, only the truncated phosphopantetheine arm of CoA is shown. **B–D** Biosensor output of the individual HvCHS2 variant libraries, measured 24 h after supplementation with 0.2 mM ferulic acid (**B**), 0.5 mM 4-methoxycinnamic acid (**C**), or 0.5 mM isoferulic acid (**D**). The median biosensor output is indicated by a black line, with lower and upper quartiles are shown in blue. The median biosensor output of the parent enzyme is indicated by a dotted line (*n* = 96). **E** Heat map showing fold-change in absolute product titres for each enzyme variant relative to the parent enzyme. Absolute titres are mean values of three biological replicates. **F–I** Correlation between product titres and biosensor output for HvCHS2 variants, measured 24 h after feeding with 0.5 mM 4-methoxycinnamic acid (**F**), 0.5 mM isoferulic acid (**G**), 0.5 mM ferulic acid (**H**), and 1 mM ferulic acid (**I**). Each data point represents the mean, with error bars indicating the standard deviation of three biological replicates, except for the parent enzyme, which is represented by the mean ± standard deviation of six biological replicates. The Pearson correlation coefficient r^2^ is shown (*n* = 22), together with the 95% confidence interval of the linear regression.

Individual site-saturation libraries were generated for each of the seventeen selected residues using HvCHS2 carrying the Q232P and D234V substitutions as the mutational template, rather than the wild-type enzyme, and cloned into a plasmid pathway background comprising Gm4CL1, MsCHI1, and the FdeR-based biosensor. The resulting mutant libraries were transformed into *E. coli* strain SBC016072, and 96 individual clones per library were screened following supplementation with ferulic acid, isoferulic acid, or 4-methoxycinnamic acid. Biosensor output was quantified 24 h after substrate addition, and variants exhibiting reporter activity comparable to or exceeding that of the parent HvCHS2 variant with at least one substrate were selected for sequencing.

Across all three substrates tested, mutations at the surface-exposed residues C204 and D317 had little to no effect on biosensor output compared to the parent enzyme, indicating a limited impact on product formation (Figure 5B–D). In contrast, mutations within the active-site cavity and CoA-binding tunnel resulted in diverse biosensor output profiles, reflecting varying sensitivities of individual positions to amino acid substitution. Certain residues, such as Q163 and T196, exhibited substantial mutational plasticity, as substitutions at these positions yielded variants with reporter outputs ranging from low to high levels. By contrast, residues including E194 and D219 appeared to be highly constrained, as substitutions at these positions consistently led to diminished biosensor output. Inspection of the active site of HvCHS2 complexed with CoA and eriodictyol shows that the side chains of E194 and D219 are central to a hydrogen bond network that includes interactions both with other key active site residues and the phenol groups of eriodictyol, and therefore likely play a large part in orchestrating substrate acceptance for several of the ring-substituted cinnamoyl-CoA derivatives.

Sequenced variants selected based on biosensor output were retransformed into *E. coli* strain SBC016072, and flavanone production was quantified analytically following supplementation with 0.5 mM ferulic acid, isoferulic acid, or 4-methoxycinnamic acid, as well as with 1 mM ferulic acid. Several variants exhibited statistically significant improvements in homoeriodictyol production relative to the parent HvCHS2 enzyme (Figure 5E). At 0.5 mM ferulic acid, variants I195S, I195C, T196S, and M339C showed increased product titres, with I195C yielding 94 mg/L homoeriodictyol, corresponding to a 20% improvement over the parent enzyme (Figure S3). At 1 mM ferulic acid, six variants (I195C, C204S, C204D, M339V, M339C, and M339T) outperformed the parent enzyme, with M339C reaching 161 mg/L homoeriodictyol, representing a 15% increase (Figure S4).

Comparison of homoeriodictyol production at the two ferulic acid concentrations revealed broadly similar conversion trends for most variants. However, T196S and S340A deviated from this trend, exhibiting higher or comparable homoeriodictyol titres relative to the parent enzyme at 0.5 mM ferulic acid, but reduced homoeriodictyol titres at 1 mM. This behaviour may indicate increased substrate inhibition by feruloyl-CoA in these variants, leading to reduced performance at higher substrate concentrations^39^.

Enhanced production of *para*-*O*-methylated flavanones was also observed. Upon supplementation with isoferulic acid, variants M339V, M339C, M339I, and S340I accumulated significantly higher hesperetin titres than the parent enzyme, with M339V yielding 9.1 mg/L hesperetin, corresponding to an approximately twofold improvement (Figure S5). Similarly, feeding with 4-methoxycinnamic acid resulted in increased isosakuranetin production for variants C204G, M339V, and M339I, with M339V reaching 10.5 mg/L isosakuranetin, representing an over twofold increase relative to the parent enzyme (Figure S6).

Finally, we assessed the extent to which biosensor output correlated with analytically quantified product titres across enzyme variants and substrates. For isosakuranetin, biosensor output showed a strong positive correlation with product titre, indicating that FdeR-dependent reporter activation provides a reliable proxy for product formation under these conditions (Figure 5F). In contrast, no significant correlation was observed for hesperetin, suggesting that product titres are not within the ligand detection range of the biosensor (Figure 5G). The presence of outliers exhibiting high biosensor output but low hesperetin titres (e.g. variant T196G) suggests that certain HvCHS2 variants may give rise to alternative metabolites capable of activating the biosensor. For homoeriodictyol, biosensor output correlated well with product titre at both ferulic acid concentrations tested, albeit with increased variability at higher substrate levels (Figure 5H and 5I). This increased variability may reflect partial saturation of the biosensor at higher substrate concentrations. Notably, variant S340A deviated from this trend at 0.5 mM ferulic acid, yielding homoeriodictyol levels comparable to the parent enzyme but a considerably lower biosensor output. This observation may reflect reduced apparent turnover or delayed product formation, potentially consistent with mild substrate inhibition, resulting in lower intracellular accumulation of the activating ligand over time and consequently reduced activation of the biosensor and reporter gene expression. Despite individual outliers, these results demonstrate that the FdeR-based biosensor provides a robust screening tool when product titres fall within its effective detection range, supporting its application in enzyme engineering workflows.

### 3.4 Variant-dependent formation of isoferuloyllderived derailment products by HvCHS2

During the analysis of HvCHS2 variants within the three-enzyme pathway comprising Gm4CL1, HvCHS2, and MsCHI1, two variants in particular exhibited behaviour distinct from the remainder of the variant set. Relative to the parent HvCHS2 enzyme, substitutions S340I and S340L resulted in a more than threefold reduction in residual isoferulic acid levels that was not accompanied by a proportional increase in hesperetin production (FigureD6A, Figure S5). UPLC-DAD analysis of culture extracts from strains expressing these variants revealed the appearance of two prominent peaks that were present only at low levels in the parent enzyme background, indicating a shift in product profile and prompting further investigation into the chemical identity of these species.

As a typeDIII polyketide synthase, chalcone synthase is known to generate derailment products when control over polyketide chain elongation or the subsequent regiospecific intramolecular Claisen condensation is disrupted. Such derailment can result in premature hydrolysis or non-enzymatic lactonisation of polyketide intermediates, yielding truncated or alternative products^22, 24^. For example, incomplete extension of a *p*-coumaroyl-derived polyketide leads to release of the triketide intermediate as bisnoryangonin (BNY), whereas failure to catalyse the Claisen condensation at the tetraketide stage diverts the pathway toward formation of *p*-coumaroyl-triacetic acid lactone (CTAL)^23^.

Similar behaviour has been reported for chalcone synthase from *M. sativa*, where mutation of the active-site residue G256 resulted in the formation of corresponding triketide- and tetraketide-derived derailment products from a range of starter molecules, including feruloyl-CoA^23^. Based on these observations, we hypothesised that HvCHS2 variants S340I and S340L likewise promote the formation of isoferuloyl-derived derailment products (FigureD6B).

By analogy to previously reported CHS derailment products, the observed species are likely to be (*E*)-4-hydroxy-6-(4-(3-hydroxy-4-methoxyphenyl)-2-oxobut-3-en-1-yl)-2*H*-pyran-2-one, arising from derailment at the tetraketide stage (hereafter termed derailment product A), and (*E*)-4-hydroxy- 6-(3-hydroxy-4-methoxystyryl)-2*H*-pyran-2-one, derived from the triketide intermediate (hereafter termed derailment product B). In addition, formation of 6-methyl-4-hydroxy-2-pyrone (methylpyrone), which can arise independently of the isoferuloyl-CoA starter unit, from CHS activity on malonyl-CoA alone, cannot be excluded^23, 40^.

Although authentic standards for derailment productsDA andDB are not commercially available, LC-QTof MS analysis provided multiple lines of supporting evidence for their assignment. Accurate-mass measurements of the precursor ions corresponding to peaksDA andDB (Figure 6A) were in good agreement with the calculated masses for C_16_H_14_O_6_ and C_14_H_12_O_5_, respectively, with relative mass errors below 3 ppm. Furthermore, MS^E^ analysis revealed diagnostic product ions for both peaks that were consistent with those reported previously for tetraketide- and triketide-derived derailment products formed from feruloyl-CoA^23^, providing additional support for these assignments. By contrast, although formation of methylpyrone might have occured, its presence could not be unambiguously confirmed based on the available accurate-mass and fragmentation data.

**Figure 6.**
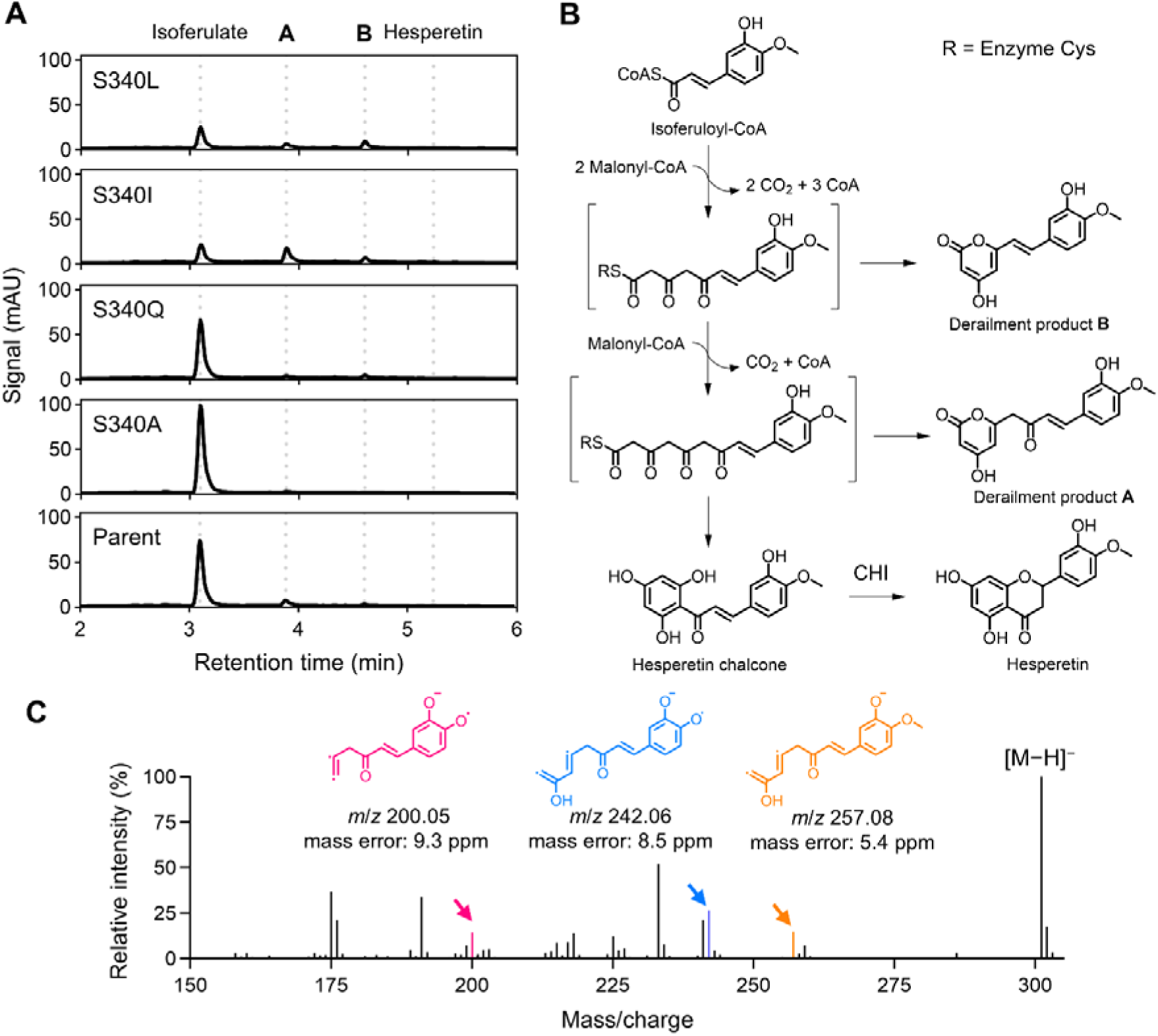
Biosynthesis of isoferuloyl-derived derailment products by HvCHS2. **A** Representative UPLC-DAD chromatograms (290 nm) of culture extracts from the parent HvCHS2 strain and the S340L, S340I, S340Q, and S340A variants. Dashed lines indicate the retention times of isoferulic acid, hesperetin, and derailment products A and B. **B** Reaction pathway leading from isoferuloyl-CoA to the canonical CHS product hesperetin chalcone and to derailment products A and B via tetraketide-and triketide-based intermediates. **C** MS^E^ ESI^−^ mass spectrum (15–40 eV collision-energy ramp) extracted at the retention time corresponding to peak A observed in the HvCHS2 S340I variant. Selected product ion structures, including [M−H−CO_2_]^−^ (orange), [M−H−CO_2_−CH_3_]^−^ (blue), and [M−H−CO_2_−CH_3_−C_2_H_2_O]^−^ (magenta) are assigned to their corresponding *m*/*z* signals.

Because derailment product A and the canonical CHS product hesperetin chalcone have identical molecular masses, the observed MS^E^ fragmentation data for peak A were compared with those predicted for both species. Diagnostic product ions consistent with derailment product A were detected (Figure 6C), whereas no product ions uniquely attributable to hesperetin chalcone were observed. Moreover, even if hesperetin chalcone were to accumulate due to incomplete cyclisation in the subsequent CHI-catalysed step, spontaneous cyclisation would be expected to yield an enantiomeric mixture of hesperetin^41, 42^, making it unlikely that peak A corresponds to hesperetin chalcone. Together, these data support the assignment of peaks A and B to isoferuloyl-derived derailment products A and B, respectively, rather than to canonical intermediates of hesperetin biosynthesis.

Analysis of extracted-ion chromatograms corresponding to the deprotonated precursor ions [M−H]^−^ of derailment products A and B across the analysed HvCHS2 variants showed signal intensities that closely mirrored those observed in the UPLC-DAD chromatograms at 290Dnm (Figures S7, S8, and Figure 6A). Among the variants analysed, HvCHS2 S340I produced the highest level of derailment product A, whereas S340L exhibited the highest accumulation of derailment product B, followed by S340I. Lower levels of both derailment products were also detected in strains expressing HvCHS2 S340A, S340Q, and the parent HvCHS2 variant, indicating that product derailment is not exclusive to the engineered variants but is enhanced by specific substitutions at this position.

Collectively, these results indicate that substitution of residue S340 in HvCHS2 with isoleucine or leucine promotes diversion of isoferuloyl-derived intermediates toward triketide- and tetraketide-derived derailment products. A similar effect has been reported for mutation of the corresponding residue in *M. sativa* CHS (S338I), which led to enhanced formation of *p*-coumaroyl-derived derailment products such as BNY and CTAL^22^, underscoring the importance of this position in governing chalcone formation. While these substitutions only modestly (in the case of S340I) or not at all (in the case of S340L) enhance hesperetin biosynthesis, they may nevertheless provide a basis for the targeted bioproduction of defined polyketide derailment products.

In plants, association of CHS with chalcone isomerase-like proteins has been shown to suppress derailment and promote efficient flavanone formation^43, 44^. Introduction of such accessory proteins has similarly been used in *E. coli* to reduce CTAL formation and enhance naringenin production^45^. Given that low-level derailment products were also detected for the parent HvCHS2 variant in this study, co-expression of a chalcone isomerase-like protein represents a potential strategy to further suppress derailment and increase hesperetin titres.

### 3.5 Directed evolution of AtCHS enhances production of fluoro-substituted flavanones

Building on the successful application of the FdeR-based biosensor for the directed evolution of HvCHS2, we next applied the same screening strategy to engineer *A. thaliana* chalcone synthase (AtCHS). The biosensor was used to identify AtCHS variants with enhanced capacity to support the biosynthesis of fluoro-substituted flavanones, specifically 3′-fluoro-5,7-dihydroxyflavanone and 4′-fluoro-5,7-dihydroxyflavanone. Enhancement of 2′-fluoro-5,7-dihydroxyflavanone production was not pursued further, as high conversion yields and product titres were already achieved with the wild-type enzyme under the conditions tested.

Based on the HvCHS2 mutational landscape, residues I195, T196, and M339 were identified as positions exhibiting mutational plasticity and yielding variants with improved biosynthesis of *O*-methylated flavanones. The corresponding residues in AtCHS (I198, T199, and M343) were therefore selected for site-saturation mutagenesis.

Individual site-saturation libraries were generated for each of the three selected residues and cloned into plasmid SBC016443, comprising Gm4CL4, MsCHI1, and the FdeR-based biosensor. The resulting libraries were transformed into *E. coli* strain SBC016072, and 96 individual clones per library were screened following supplementation with either 0.5 mM 3-fluorocinnamic acid or 0.5 mM 4-fluorocinnamic acid. Biosensor output was quantified 24 h after substrate addition, and variants exhibiting reporter activity comparable to or exceeding that of the wildDtype AtCHS enzyme for at least one substrate were selected for sequencing.

Analysis of biosensor output revealed that substitutions at residues I198 and T199 were generally better tolerated than those at M343 under both substrate conditions (Figure 7A and 7B). In contrast, substitutions at M343 consistently resulted in considerable reduced reporter output relative to the wild-type enzyme, indicating that perturbation at this position adversely affects conversion of 3-, and 4-fluorocinnamoyl-CoA into the corresponding chalcones.

**Figure 7.**
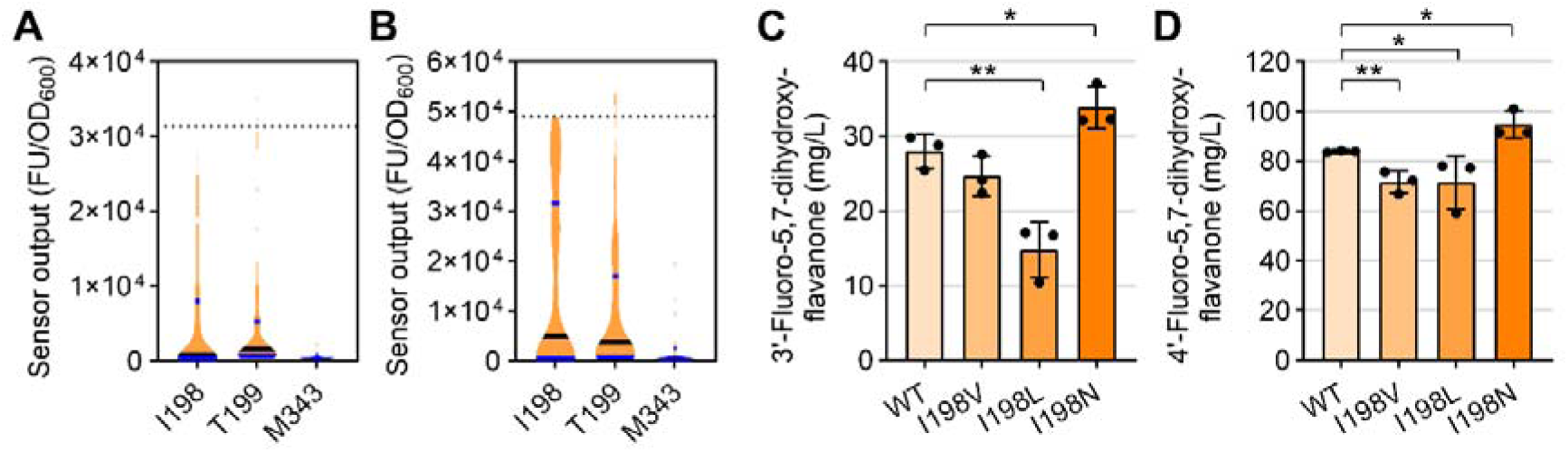
Directed evolution of AtCHS. **A** and **B** Biosensor output of AtCHS variant libraries, measured 24 h after supplementation with 0.5 mM 3-fluorocinnamic acid (**A**) or 4-fluorocinnamic acid (**B**). The median biosensor output is indicated by a black line, with lower and upper quartiles shown in blue (*n* = 96). The median biosensor output of the wild-type enzyme is indicated by a dotted line. **C** and **D** Absolute titres of 3′-fluoro-5,7-dihydroxyflavanone (**C**) and 4′-fluoro-5,7-dihydroxyflavanone (**D**) for the I198 variants and the wild-type enzyme (WT). Data are presented as mean ± standard deviation (*n* = 3). Statistical significance is indicated as * where *p* < 0.05, and ** where *p* < 0.01 (unpaired two-tailed *t* test).

Sequencing of selected clones identified three substitutions at residue I198 – namely I198V, I198L, and I198N – that yielded biosensor outputs comparable to or exceeding that of the wild-type enzyme in response to feeding with at least one of the two fluoro-substituted cinnamic acid substrates. These variants were subsequently retransformed into *E. coli* strain SBC016072, and flavanone production was quantified analytically following supplementation with 0.5 mM 3-fluorocinnamic acid or 0.5 mM 4-fluorocinnamic acid.

Of the three I198 variants tested, only I198N demonstrated improved performance relative to the wild-type enzyme for both substrates. This variant produced 34 mg/L 3′-fluoro-5,7-dihydroxyflavanone and 95 mg/L 4′-fluoro-5,7-dihydroxyflavanone, corresponding to statistically significant increases in product titres of 21% and 13%, respectively (Figure 7C and 7D).

Similar to residue I195 in HvCHS2, I198 in AtCHS is located on a loop between secondary structural features, and substitution at this position (I198N) likely alters local packing or conformational flexibility within the active site, thereby leading to improved accommodation of 3-and 4-fluorocinnamoyl-CoA substrates.

## 4. Conclusion

In this study, we integrated combinatorial pathway optimisation with biosensor-guided enzyme engineering to enhance the microbial production of structurally diverse flavanones. By systematically evaluating combinations of 4CL, CHS, and CHI, we identified pathway configurations that enable efficient conversion of ring-substituted cinnamic acids into both natural and non-natural flavanones. Analysis of pathway performance revealed a substrate-dependent shift in pathway control, with CHS representing the primary bottleneck for the conversion of bulkier or methoxy-substituted substrates.

In parallel, we established an FdeR-based biosensor as a broadly responsive and versatile tool for monitoring flavanone biosynthesis and enabling high-throughput screening. Using this platform, we performed directed evolution of chalcone synthases from *H. vulgare* and *A. thaliana*, identifying variants that enhance production of *O*-methylated and fluoro-substituted flavanones, as well as variants that promote the formation of isoferuloyl-derived derailment products.

Collectively, this work demonstrates that early-stage diversification through exploitation of pathway promiscuity provides an effective route to access structurally diverse flavanones without reliance on downstream tailoring enzymes. Furthermore, integration of biosensor-guided screening with combinatorial pathway design establishes a generalisable framework for the rapid optimisation of biosynthetic pathways and enzyme engineering.

These findings provide a foundation for expanding flavonoid diversity in microbial systems and highlight opportunities to further improve pathway performance through enzyme engineering and biosensor tuning, enabling access to increasingly complex and non-natural phenylpropanoid-derived compounds.

## Acknowledgements

This work was supported by the Future Biomanufacturing Research Hub (grant EP/S01778X/1), funded by the Engineering and Physical Sciences Research Council (EPSRC) and Biotechnology and Biological Sciences Research Council (BBSRC) as part of UK Research and Innovation (UKRI). The authors thank the Faculty of Science and Engineering Mass Spectrometry and Separation Facility (RRID:DSCR_024761) and Technical Specialist for Separations, Dr Maira Hernandez-Guzman.

## Author contributions: CRediT

**EKRH:** Conceptualisation, Formal analysis, Investigation, Methodology, Visualisation, Writing – original draft. **CK:** Formal analysis, Investigation, Methodology, Visualisation, Writing – review and editing. **JNW:** Formal analysis, Investigation, Visualisation, Writing – review and editing. **RS:** Formal analysis, Investigation, Methodology, Writing – review and editing. **CJR:** Resources, Writing – review and editing. **NSS:** Funding acquisition, Writing – review and editing.

## Data availability

Plasmids SBC016302, SBC016397, SBC016420, SBC016421, SBC016434, SBC016441, SBC016442, and SBC016443 are available from Addgene under accession numbers 258001–258005 and 258007–258009, respectively.

## Appendices

Appendix A: Supplementary Information

## References

[1.] Williams, C. A., and Grayer, R. J. (2004) Anthocyanins and other flavonoids, Nat. Prod. Rep. 21, 539–573. 10.1039/B311404J.

[2.] Chen, S., Wang, X., Cheng, Y., Gao, H., and Chen, X. (2023) A review of classification, biosynthesis, biological activities and potential applications of flavonoids, Molecules 28, 4982. 10.3390/molecules28134982.

[3.] Mierziak, J., Kostyn, K., and Kulma, A. (2014) Flavonoids as important molecules of plant interactions with the environment, Molecules 19, 16240–16265. 10.3390/molecules191016240.

[4.] Câmara, J. S., Locatelli, M., Pereira, J. A., Oliveira, H., Arlorio, M., Fernandes, I., Perestrelo, R., Freitas, V., and Bordiga, M. (2022) Behind the scenes of anthocyanins—From the health benefits to potential applications in food, pharmaceutical and cosmetic fields, Nutrients 14, 5133. 10.3390/nu14235133.

[5.] Dias, M. C., Pinto, D. C., and Silva, A. M. (2021) Plant flavonoids: Chemical characteristics and biological activity, Molecules 26, 5377. 10.3390/molecules26175377.

[6.] Stachelska, M. A., Karpiński, P., and Kruszewski, B. (2025) A comprehensive review of biological properties of flavonoids and their role in the prevention of metabolic, cancer and neurodegenerative diseases, Appl. Sci. 15, 10840. 10.3390/app151910840.

[7.] Carbonell, P., Jervis, A. J., Robinson, C. J., Yan, C., Dunstan, M., Swainston, N., Vinaixa, M., Hollywood, K. A., Currin, A., Rattray, N. J., Taylor, S., Spiess, R., Sung, R., Williams, A. R., Fellows, D., Stanford, N. J., Mulherin, P., Le Feuvre, R., Barran, P., Goodacre, R., Turner, N. J., Goble, C., Guoqiang Chen, G., Kell, D. B., Micklefield, J., Breitling, R., Takano, E., Faulon, J.-L., and Scrutton, N. S. (2018) An automated Design-Build-Test-Learn pipeline for enhanced microbial production of fine chemicals, *Commun*. Biol. 1, 1–10. 10.1038/s42003-018-0076-9.

[8.] Trantas, E. A., Koffas, M. A., Xu, P., and Ververidis, F. (2015) When plants produce not enough or at all: metabolic engineering of flavonoids in microbial hosts, Front. Plant Sci. 6, 7. 10.3389/fpls.2015.00007.

[9.] Chaves, J. O., De Souza, M. C., Da Silva, L. C., Lachos-Perez, D., Torres-Mayanga, P. C., Machado, A. P. d. F., Forster-Carneiro, T., Vázquez-Espinosa, M., González-de-Peredo, A. V., Barbero, G. F., and Rostagno, M. A. (2020) Extraction of flavonoids from natural sources using modern techniques, Front. Chem. 8, 507887. 10.3389/fchem.2020.507887.

[10.] Zhang, Q., Yu, S., Lyu, Y., Zeng, W., and Zhou, J. (2021) Systematically engineered fatty acid catabolite pathway for the production of (2*S*)-naringenin in *Saccharomyces cerevisiae*, ACS Synth. Biol. 10, 1166–1175. 10.1021/acssynbio.1c00002.

[11.] Tong, Y., Li, N., Zhou, S., Zhang, L., Xu, S., and Zhou, J. (2024) Improvement of chalcone synthase activity and high-efficiency fermentative production of (2*S*)-naringenin via *in vivo* biosensor-guided directed evolution, ACS Synth. Biol. 13, 1454–1466. 10.1021/acssynbio.3c00570.

[12.] Wu, J., Zhang, X., Zhou, J., and Dong, M. (2016) Efficient biosynthesis of (2S)-pinocembrin from D-glucose by integrating engineering central metabolic pathways with a pH-shift control strategy, Bioresour. Technol. 218, 999–1007. 10.1016/j.biortech.2016.07.066.

[13.] Qiu, Z., Han, Y., Li, J., Ren, Y., Liu, X., Li, S., Zhao, G.-R., and Du, L. (2025) Metabolic division engineering of Escherichia coli consortia for de novo biosynthesis of flavonoids and flavonoid glycosides, Metab. Eng. 89, 60–75. 10.1016/j.ymben.2025.02.001.

[14.] Kim, B.-G., Lee, Y., Hur, H.-G., Lim, Y., and Ahn, J.-H. (2006) Flavonoid 3′-*O*-methyltransferase from rice: cDNA cloning, characterization and functional expression, Phytochemistry 67, 387–394. 10.1016/j.phytochem.2005.11.022.

[15.] Lee, H., Park, S., Lee, S. B., Song, J., Kim, T.-H., and Kim, B.-G. (2024) Tailored biosynthesis of diosmin through reconstitution of the flavonoid pathway in *Nicotiana benthamiana*, Front. Plant Sci. 15, 1464877. 10.3389/fpls.2024.1464877.

[16.] Hugueney, P., Provenzano, S., Verriès, C., Ferrandino, A., Meudec, E., Batelli, G., Merdinoglu, D., Cheynier, V., Schubert, A., and Ageorges, A. (2009) A novel cation-dependent *O*-methyltransferase involved in anthocyanin methylation in grapevine, Plant Physiol. 150, 2057–2070. 10.1104/pp.109.140376.

[17.] Abe, I., Morita, H., Nomura, A., and Noguchi, H. (2000) Substrate specificity of chalcone synthase: enzymatic formation of unnatural polyketides from synthetic cinnamoyl-CoA analogues, J. Am. Chem. Soc. 122, 11242–11243. 10.1021/ja0027113.

[18.] Kufs, J. E., Hoefgen, S., Rautschek, J., Bissell, A. U., Graf, C., Fiedler, J., Braga, D., Regestein, L., Rosenbaum, M. A., Thiele, J., and Valiante, V. (2020) Rational design of flavonoid production routes using combinatorial and precursor-directed biosynthesis, ACS Synth. Biol. 9, 1823–1832. 10.1021/acssynbio.0c00172.

[19.] Kubiak, J., Szyk, P., Czarczynska-Goslinska, B., and Goslinski, T. (2025) Flavonoids, Chalcones, and Their Fluorinated Derivatives—Recent Advances in Synthesis and Potential Medical Applications, Molecules 30, 2395. 10.3390/molecules30112395.

[20.] Austin, M. B., and Noel, J. P. (2003) The chalcone synthase superfamily of type III polyketide synthases, Nat. Prod. Rep. 20, 79–110. 10.1039/B100917F.

[21.] Ortiz, A., and Sansinenea, E. (2023) Phenylpropanoid Derivatives and Their Role in Plants’ Health and as antimicrobials, Curr. Microbiol. 80, 380. 10.1007/s00284-023-03502-x.

[22.] Jez, J. M., Austin, M. B., Ferrer, J.-L., Bowman, M. E., Schröder, J., and Noel, J. P. (2000) Structural control of polyketide formation in plant-specific polyketide synthases, Chem. Biol. 7, 919–930. 10.1016/S1074-5521(00)00041-7.

[23.] Jez, J. M., Bowman, M. E., and Noel, J. P. (2001) Structure-guided programming of polyketide chain-length determination in chalcone synthase, Biochemistry 40, 14829–14838. 10.1021/bi015621z.

[24.] Jez, J., Ferrer, J., Bowman, M., Austin, M., Schröder, J., Dixon, R., and Noel, J. (2001) Structure and mechanism of chalcone synthase-like polyketide synthases, J. Ind. Microbiol. Biotechnol. 27, 393–398. 10.1038/sj.jim.7000188.

[25.] Hwang, H. G., Milito, A., Yang, J.-S., Jang, S., and Jung, G. Y. (2023) Riboswitch-guided chalcone synthase engineering and metabolic flux optimization for enhanced production of flavonoids, Metab. Eng. 75, 143–152. 10.1016/j.ymben.2022.12.006.

[26.] Alguel, Y., Meng, C., Terán, W., Krell, T., Ramos, J. L., Gallegos, M.-T., and Zhang, X. (2007) Crystal structures of multidrug binding protein TtgR in complex with antibiotics and plant antimicrobials, J. Mol. Biol. 369, 829–840. 10.1016/j.jmb.2007.03.062.

[27.] Wassem, R., Marin, A., Daddaoua, A., Monteiro, R., Chubatsu, L., Ramos, J., Deakin, W., Broughton, W., Pedrosa, F., and Souza, E. (2017) A NodDDlike protein activates transcription of genes involved with naringenin degradation in a flavonoidDdependent manner in *Herbaspirillum seropedicae*, Environ. Microbiol. 19, 1030–1040. doi.org/10.1111/1462-2920.13604.

[28.] Hanko, E. K., Robinson, C. J., Bhanot, S., Jervis, A. J., and Scrutton, N. S. (2024) Engineering an *Escherichia coli* strain for enhanced production of flavonoids derived from pinocembrin, Microb. Cell Fact. 23, 312. 10.1186/s12934-024-02582-z.

[29.] Swainston, N., Dunstan, M., Jervis, A. J., Robinson, C. J., Carbonell, P., Williams, A. R., Faulon, J.-L., Scrutton, N. S., and Kell, D. B. (2018) PartsGenie: an integrated tool for optimizing and sharing synthetic biology parts, Bioinformatics 34, 2327–2329. 10.1093/bioinformatics/bty105.

[30.] Sambrook, J., and Russell, D. W. (2001) Molecular Cloning: A Laboratory Manual, 3rd ed., Cold Spring Harbor Laboratory Press, New York.

[31.] Wang, F., Allen, D., Tian, S., Oler, E., Gautam, V., Greiner, R., Metz, T. O., and Wishart, D. S. (2022) CFM-ID 4.0 – a web server for accurate MS-based metabolite identification, Nucleic Acids Res. 50, W165–W174. 10.1093/nar/gkac383.

[32.] Katsuyama, Y., Funa, N., Miyahisa, I., and Horinouchi, S. (2007) Synthesis of unnatural flavonoids and stilbenes by exploiting the plant biosynthetic pathway in *Escherichia coli*, Chem. Biol. 14, 613–621. 10.1016/j.chembiol.2007.05.004.

[33.] Cui, H., Song, M. C., Lee, J. Y., and Yoon, Y. J. (2019) Microbial production of *O*-methylated flavanones from methylated phenylpropanoic acids in engineered *Escherichia coli*, J. Ind. Microbiol. Biotechnol. 46, 1707–1713. 10.1007/s10295-019-02239-6.

[34.] Peng, B., Zhang, L., He, S., Oerlemans, R., Quax, W. J., Groves, M. R., and Haslinger, K. (2023) Engineering a plant polyketide synthase for the biosynthesis of methylated flavonoids, J. Agric. Food Chem. 72, 529–539. 10.1021/acs.jafc.3c06785.

[35.] Dunstan, M. S., Robinson, C. J., Jervis, A. J., Yan, C., Carbonell, P., Hollywood, K. A., Currin, A., Swainston, N., Feuvre, R. L., Micklefield, J., Faulon, J.-L., Breitling, R., Turner, N. J., Takano, E., and Scrutton, N. S. (2020) Engineering Escherichia coli towards de novo production of gatekeeper (2S)-flavanones: naringenin, pinocembrin, eriodictyol and homoeriodictyol, Synth. Biol. 5, ysaa012. 10.1093/synbio/ysaa012.

[36.] Knobloch, K. H., and Hahlbrock, K. (1975) Isoenzymes of *p*Dcoumarate: CoA ligase from cell suspension cultures of *Glycine max*, Eur. J. Biochem. 52, 311–320. 10.1111/j.1432-1033.1975.tb03999.x.

[37.] Lindermayr, C., Möllers, B., Fliegmann, J., Uhlmann, A., Lottspeich, F., Meimberg, H., and Ebel, J. (2002) Divergent members of a soybean (*Glycine max* L.) 4Dcoumarate: coenzyme A ligase gene family: Primary structures, catalytic properties, and differential expression, Eur. J. Biochem. 269, 1304–1315. 10.1046/j.1432-1033.2002.02775.x.

[38.] Chemler, J. A., Yan, Y., Leonard, E., and Koffas, M. A. (2007) Combinatorial mutasynthesis of flavonoid analogues from acrylic acids in microorganisms, Org. Lett. 9, 1855–1858. 10.1021/ol0703736.

[39.] Zhao, S., Jones, J. A., Lachance, D. M., Bhan, N., Khalidi, O., Venkataraman, S., Wang, Z., and Koffas, M. A. (2015) Improvement of catechin production in *Escherichia coli* through combinatorial metabolic engineering, Metab. Eng. 28, 43–53. 10.1016/j.ymben.2014.12.002.

[40.] Eckermann, S., Schröder, G., Schmidt, J., Strack, D., Edrada, R. A., Helariutta, Y., Elomaa, P., Kotilainen, M., Kilpeläinen, I., Proksch, P., Teeri, T. H., and Schröder, J. (1998) New pathway to polyketides in plants, Nature 396, 387–390. 10.1038/24652.

[41.] Jez, J. M., Bowman, M. E., Dixon, R. A., and Noel, J. P. (2000) Structure and mechanism of the evolutionarily unique plant enzyme chalcone isomerase, Nat. Struct. Biol. 7, 786–791. 10.1038/79025.

[42.] Khan, M. K., Rakotomanomana, N., Dufour, C., and Dangles, O. (2011) Binding of citrus flavanones and their glucuronides and chalcones to human serum albumin, Food Funct. 2, 617–626. 10.1039/C1FO10077G.

[43.] Waki, T., Mameda, R., Nakano, T., Yamada, S., Terashita, M., Ito, K., Tenma, N., Li, Y., Fujino, N., Uno, K., Yamashita, S., Aoki, Y., Denessiouk, K., Kawai, Y., Sugawara, S., Saito, K., Yonekura-Sakakibara, K., Morita, Y., Hoshino, A., Takahashi, S., and Nakayama, T. (2020) A conserved strategy of chalcone isomerase-like protein to rectify promiscuous chalcone synthase specificity, Nat. Commun. 11, 870. 10.1038/s41467-020-14558-9.

[44.] Ban, Z., Qin, H., Mitchell, A. J., Liu, B., Zhang, F., Weng, J.-K., Dixon, R. A., and Wang, G. (2018) Noncatalytic chalcone isomerase-fold proteins in *Humulus lupulus* are auxiliary components in prenylated flavonoid biosynthesis, Proc. Natl. Acad. Sci. U. S. A. 115, E5223–E5232. 10.1073/pnas.1802223115.

[45.] Liu, X., Li, L., and Zhao, G.-R. (2022) Systems metabolic engineering of Escherichia coli coculture for de novo production of genistein, ACS Synth. Biol. 11, 1746–1757. 10.1021/acssynbio.1c00590.

