## Supplementary Information for "Biosensor-guided evolution of chalcone synthase enhances biosynthesis of natural and non-natural flavanones"

### Supplementary Methods

#### Plasmid construction

Plasmid SBC016239 was constructed by restriction enzyme cloning. The gBlock FdeR (Table S3) was digested with AatII and NdeI and ligated into pBbE1c-rfp digested with the same enzymes.

Plasmid SBC016302 was constructed by HiFi DNA Assembly from six parts. The first part, containing the FdeR-based biosensor linked to the *rfp* reporter gene, was amplified by PCR from plasmid SBC016239 using primers EHfbrh_081 and EHfbrh_215. The second part, containing the tetracycline resistance marker, was amplified by PCR from pCAT207 using primers EHfbrh_196 and EHfbrh_197. The third part, containing the *lacI* gene and CHI2 from *Arabidopsis thaliana*, was amplified by PCR from plasmid SBC006845 using primers EHfbrh_216 and EHfbrh_217. The fourth part, containing 4CL4 from *Glycine max*, was amplified by PCR from plasmid SBC006845 using primers EHfbrh_218 and EHfbrh_219. The fifth part, containing the *trc* promoter, was amplified by PCR from plasmid SBC006845 using primers EHfbrh_220 and EHfbrh_221. The sixth part, containing CHS from *A. thaliana*, the ColE1 origin of replication, and the kanamycin resistance marker, was amplified by PCR from plasmid SBC006845 using primers EHfbrh_222 and EHfbrh_223.

Plasmid SBC016397 was constructed by HiFi DNA Assembly from two parts. The first part, containing 4CL1 from *G. max*, was amplified by PCR from gBlock Gm4CL1 using primers EHfbrh_224 and EHfbrh_284. The second part consisted of PacI- and AscI-digested plasmid SBC016302, into which the amplified part was assembled.

Plasmid SBC016420 was constructed by HiFi DNA Assembly from two parts. The first part, containing CHS2 from *Hordeum vulgare*, was amplified by PCR from gBlock HvCHS2 using primers EHfbrh_288 and EHfbrh_289. The second part consisted of SbfI- and XhoI-digested plasmid SBC016397, into which the amplified part was assembled.

Plasmid SBC016421 was constructed by HiFi DNA Assembly from two parts. The first part, containing CHI1 from *Medicago sativa*, was amplified by PCR from gBlock MsCHI using primers EHfbrh_292 and EHfbrh_293. The second part, containing 4CL1 from *G. max*, CHS2 from *H. vulgare*, the FdeR-based biosensor linked to the *rfp* reporter gene, the tetracycline resistance marker, the *lacI* gene, the ColE1 origin of replication, and the kanamycin resistance marker, was amplified by PCR from plasmid SBC016420 using primers EHfbrh_290 and EHfbrh_291.

Plasmid SBC016434 was constructed by restriction enzyme cloning. Plasmid SBC016302 was digested with SbfI and XhoI and the fragment containing *A. thaliana* CHS was ligated into plasmid SBC016421 digested with the same enzymes.

Plasmid SBC016441 was constructed by restriction enzyme cloning. Plasmid SBC016421 was digested with SbfI and XhoI and the fragment containing *H. vulgare* CHS2 was ligated into plasmid SBC016302 digested with the same enzymes.

Plasmid SBC016442 was constructed by restriction enzyme cloning. Plasmid SBC016302 was digested with PacI and AscI and the fragment containing 4CL4 from *G. max* was ligated into plasmid SBC016421 digested with the same enzymes.

Plasmid SBC016443 was constructed by restriction enzyme cloning. Plasmid SBC016302 was digested with PacI and AscI and the fragment containing 4CL4 from *G. max* was ligated into plasmid SBC016434 digested with the same enzymes.

Plasmids for site saturation mutagenesis of CHS2 from *H. vulgare* were constructed by restriction enzyme cloning. HvCHS2 was first amplified by PCR in two parts. The first part was amplified using primer EHfbrh_256 and a reverse primer annealing immediately upstream of the target mutation site. The second part was amplified using primer EHfbrh_257 and a forward primer introducing the mutation via NNK degeneracy. The two parts were subsequently combined by overlap PCR, digested with SbfI and XhoI, and ligated into plasmid SBC016421 digested with the same enzymes.

Plasmids for site saturation mutagenesis of CHS from *A. thaliana* were constructed by restriction enzyme cloning. AtCHS was first amplified by PCR in two parts. The first part was amplified using primer EHfbrh_256 and a reverse primer annealing immediately upstream of the target mutation site. The second part was amplified using primer EHfbrh_257 and a forward primer introducing the mutation via NNK degeneracy. The two parts were subsequently combined by overlap PCR, digested with SbfI and XhoI, and ligated into plasmid SBC016443 digested with the same enzymes.

#### Large-scale production of flavanones

To verify compound identity and enable absolute quantification of flavanones for which authentic standards were not commercially available (5,7,2′-trihydroxyflavanone; 2′-fluoro-5,7-dihydroxyflavanone; 3′-fluoro-5,7-dihydroxyflavanone; and 4′-fluoro-5,7-dihydroxyflavanone), these compounds were produced at larger scale, purified by solid-phase extraction, and validated by nuclear magnetic resonance (NMR) spectroscopy.

For large-scale production, cultivation conditions largely mirrored those used for small-scale biosynthesis. Briefly, a single colony of *E. coli* strain SBC016072 carrying plasmid SBC016443 was used to inoculate 5 mL of production medium supplemented with kanamycin and grown at 30 °C with shaking at 200 rpm for 18 h. The resulting seed culture was used to inoculate 500 mL of production medium supplemented with kanamycin in a 2-L baffled shake flask to a starting $OD_{600}$ of 0.02. Main cultures were incubated under identical conditions. After 6 h of growth, the appropriate ring-substituted cinnamic acid substrate and IPTG were added to final concentrations of 1 mM and 0.1 mM, respectively. Cultures were incubated for a further 24 h prior to harvest.

#### Flavanone purification and verification

For solid-phase extraction (SPE) of flavanones, the culture broth (500 mL) was clarified by centrifugation at 4,000 rpm for 10 min to remove cells, followed by centrifugation of the supernatant at 14,000 rpm for 10 min to remove residual debris. The clarified supernatant was loaded onto four SPE columns (125 mL each; Phenomenex Strata-X, 33 µm polymeric reversed-phase, 200 mg/12 mL). Columns were washed twice 10 mL deionised water. Flavanones were eluted stepwise using 50%, 60%, 70%, 80%, 90%, 100% (v/v) methanol in water (2 mL per step, two fractions each), and product elution was monitored by UPLC-DAD at 290 nm. Fractions eluted with 80–100% methanol were combined, evaporated to dryness using an Eppendorf Concentrator plus, and reconstituted in 80% methanol for further purification by semi-preparative HPLC.

Semi-preparative HPLC purification was performed using an Agilent 1260 Infinity LC system equipped with an Agilent Zorbax Eclipse XBD-C18 column (9.4 × 250 mm, 5 µm) and a diode array detector set at 290 nm. Separation was achieved at room temperature under isocratic conditions at a flow rate of 5 mL/min using methanol–water mixtures of 62%, 62%, 65%, and 65% (v/v) for 3′-fluoro-5,7-dihydroxyflavanone, 4′-fluoro-5,7-dihydroxyflavanone, 5,7,2′-trihydroxyflavanone, and 2′-fluoro-5,7-dihydroxyflavanone, respectively. The injection volume was 100 µL. Collected fractions were evaporated to dryness and stored at −20 °C prior to UV-Vis and NMR analysis.

For ultraviolet-visible (UV-Vis) analysis, flavanones were resuspended in 100% methanol and spectra were recorded using an Agilent Cary 60 UV-Vis spectrophotometer at room temperature. The baseline was corrected using 100% methanol. Measurements were performed in a quartz cuvette with 10 mm path length. The molar absorptivity (ε) was determined from the slope of absorbance versus concentration over the range 10–100 μM. UV-Vis spectra of 5,7,2′-trihydroxyflavanone, 2′-fluoro-5,7-dihydroxyflavanone, 3′-fluoro-5,7-dihydroxyflavanone, and 4′-fluoro-5,7-dihydroxyflavanone are shown in Figure S9.

For structural verification by 1D- and 2D-NMR, flavanones were dissolved in acetone-$d_6$ at a concentration of 4.4–17.6 mg/mL. $^1$H, $^{13}$C, HMBC, HSQC, COSY, and NOESY NMR spectra were recorded on a Bruker Avance 500 MHz NMR spectrometer at the NMR facility of the Manchester Institute of Biotechnology. Chemical shifts (δ) are reported in ppm relative to residual solvent signals (acetone-$d_6$: $\delta_H$ 2.05 ppm, $\delta_C$ 30.6 ppm). Multiplicities are reported as s (singlet), bs (broad singlet), d (doublet), t (triplet), and m (multiplet). High-resolution mass spectrometry data were obtained using a Waters Vion IMS-QToF. Spectroscopic data for each compound are as follows:

**5,7,2′-Trihydroxyflavanone** (5,7-dihydroxy-2-(2-hydroxyphenyl)chroman-4-one).
UV (MeOH): $\lambda_{max}$ = 290 nm (ε = 14,300 L·mol$^{-1}$·cm$^{-1}$).
$^1$H-NMR (500 MHz, acetone-$d_6$): δ 12.19 (1H, s, OH-5), 9.08 (1H, bs, OH-7), 7.54 (1H, dd, *J* = 7.8, 1.2 Hz, H-3′), 7.22 (1H, dt, *J* = 7.7, 1.5 Hz, H-5′), 6.96 (1H, m, H-6′), 6.94 (1H, m, H-4′), 6.02 (1H, d, *J* = 1.9 Hz, H-8), 5.97 (1H, d, *J* = 2.0 Hz, H-6), 5.71 (1H, dd, *J* = 13.0, 2.9 Hz, H-2), 3.11 (1H, dd, *J* = 17.1, 13.0 Hz, $H_a$-3), 2.83 (1H, dd, *J* = 17.0, 3.0 Hz, $H_b$-3) ppm. One exchangeable proton (OH-2) was not observed.
$^{13}$C-NMR (125 MHz, acetone-$d_6$): δ 198.0 (C-4), 168.0 (C-8a), 165.8 (C-7), 165.4 (C-5), 155.5 (C-2′), 131.0 (C-5′), 128.5 (C-3′), 127.0 (C-1′), 121.5 (C-4′), 117.0 (C-6′), 103.9 (C-4a), 97.5 (C-6), 96.6 (C-8), 76.2 (C-2), 43.3 (C-3) ppm.
HRMS (ESI$^-$): *m/z* calculated for $C_{15}H_{11}O_5^-$, 271.06120 [M−H]$^-$; found 271.05988.
The annotated structure of 5,7,2′-trihydroxyflavanone, including key 2D-NMR correlations, is shown in Figure S10. The $^1$H- and $^{13}$C-NMR spectra are shown in Figures S11 and S12, respectively.

**2′-Fluoro-5,7-dihydroxyflavanone** (2-(2-fluorophenyl)-5,7-dihydroxy-2,3-dihydrochromen-4-one).
UV (MeOH): $\lambda_{max}$ = 290 nm (ε = 9,610 L·mol$^{-1}$·cm$^{-1}$).
$^1$H-NMR (500 MHz, acetone-$d_6$): δ 12.14 (1H, s, OH-5), 9.67 (1H, bs, OH-7), 7.71 (1H, dt, *J* = 7.6, 1.5 Hz, H-6′), 7.48 (1H, m, H-4′), 7.32 (1H, t, *J* = 7.6 Hz, H-5′), 7.22 (1H, dd, *J* = 10.7, 8.4 Hz, H-3′), 6.01 (1H, d, *J* = 1.3 Hz, H-8), 5.99 (1H, d, *J* = 1.2 Hz, H-6), 5.83 (1H, dd, *J* = 13.0, 3.0 Hz, H-2), 3.25 (1H, ddd, *J* = 17.1, 13.1,1.0 Hz, $H_a$-3), 2.81 (1H, dd, *J* = 17.1, 3.0 Hz, $H_b$-3) ppm.
$^{13}$C-NMR (125 MHz, acetone-$d_6$): δ 197.2 (C-4), 168.2 (C-7), 166.1 (C-5), 164.8 (C-8a), 161.6 (d, *J* = 246.5 Hz, C-2′), 132.3 (d, *J* = 8.5 Hz, C-4′), 129.8 (d, *J* = 3.5 Hz, C-6′), 127.6 (d, *J* = 13.1 Hz, C-1′), 126.4 (d, *J* = 3.5 Hz, C-5′), 117.2 (d, *J* = 21.7 Hz, C-3′), 103.9 (C-4a), 97.9 (C-6), 96.7 (C-8), 75.0 (d, *J* = 3.4 Hz, C-2), 43.1 (C-3) ppm. Carbon atoms at positions 2, 1′, 2′, 3′, 4′, 5′, and 6′ exhibited duplicated peaks, consistent with the influence of fluorine substitution on the aromatic B-ring[1].
HRMS (ESI$^-$): *m/z* calculated for $C_{15}H_{10}FO_4^-$, 273.05686 [M−H]$^-$; found 273.05580.
The annotated structure of 2′-fluoro-5,7-dihydroxyflavanone, including key 2D-NMR correlations, is shown in Figure S10. The $^1$H- and $^{13}$C-NMR spectra are shown in Figures S13 and S14, respectively.

**3′-Fluoro-5,7-dihydroxyflavanone** (2-(3-fluorophenyl)-5,7-dihydroxy-2,3-dihydrochromen-4-one).
UV (MeOH): $\lambda_{max}$ = 290 nm (ε = 8,380 L·mol$^{-1}$·cm$^{-1}$).
$^1$H-NMR (500 MHz, acetone-$d_6$): δ 12.13 (1H, s, OH-5), 9.72 (1H, bs, OH-7), 7.51 (1H, dt, *J* = 7.9, 6.1 Hz, H-4′), 7.41 (1H, d, *J* = 7.9 Hz, H-2′), 7.38 (1H, d, *J* = 9.9 Hz, H-6′), 7.17 (1H, dt, *J* = 8.5, 2.4 Hz, H-5′), 6.03 (1H, d, *J* = 2.0 Hz, H-8), 5.98 (1H, d, *J* = 2.1 Hz, H-6), 5.63 (1H, dd, *J* = 12.7, 3.0 Hz, H-2), 3.17 (1H, dd, *J* = 17.0, 12.7 Hz, $H_a$-3), 2.87 (1H, dd, *J* = 17.0, 3.1 Hz, $H_b$-3) ppm.
$^{13}$C-NMR (125 MHz, acetone-$d_6$): δ 197.2 (C-4), 168.1 (C-7), 165.8 (C-5), 164.7 (C-8a), 164.5 (d, *J* = 244.4 Hz, C-3′), 143.7 (d, *J* = 7.8 Hz, C-1′), 132.3 (d, *J* = 8.4 Hz, C-4′), 123.9 (d, *J* = 2.9 Hz, C-2′), 116.8 (d, *J* = 20.9 Hz, C-5′), 114.8 (d, *J* = 22.7 Hz, C-6′), 103.9 (C-4a), 97.7 (C-6), 96.7 (C-8), 79.9 (d, *J* = 5.6 Hz, C-2), 44.3 (C-3) ppm. Carbon atoms at positions 2, 1′, 2′, 3′, 4′, 5′, and 6′ exhibited duplicated peaks, consistent with the influence of fluorine substitution on the aromatic B-ring[1].

HRMS (ESI$^-$): *m/z* calculated for $C_{15}H_{10}FO_4^-$, 273.05686 $[M-H]^-$; found 273.05506.
The annotated structure of 3′-fluoro-5,7-dihydroxyflavanone, including key 2D-NMR correlations, is shown in Figure S10. The $^1$H- and $^{13}$C-NMR spectra are shown in Figures S15 and S16, respectively.

**4′-Fluoro-5,7-dihydroxyflavanone** (2-(4-fluorophenyl)-5,7-dihydroxy-2,3-dihydrochromen-4-one).
UV (MeOH): $\lambda_{max}$ = 290 nm (ε = 10,044 L·mol$^{-1}$·cm$^{-1}$).
$^1$H-NMR (500 MHz, acetone-$d_6$): δ 12.15 (1H, s, OH-5), 9.70 (1H, bs, OH-7), 7.63 (2H, dd, *J* = 8.6, 5.4 Hz, H-2′ and H-6′), 7.22 (2H, t, *J* = 8.8 Hz, H-3′ and H-5′), 6.03 (1H, d, *J* = 2.0 Hz, H-8), 5.98 (1H, d, *J* = 2.1 Hz, H-6), 5.59 (1H, dd, *J* = 12.9, 3.0 Hz, H-2), 3.17 (1H, dd, *J* = 17.0, 12.9 Hz, $H_a$-3), 2.81 (1H, dd, *J* = 17.0, 3.0 Hz, $H_b$-3) ppm.
$^{13}$C-NMR (125 MHz, acetone-$d_6$): δ 197.4 (C-4), 168.0 (C-7), 165.8 (C-5), 164.8 (C-8a), 164.4 (d, *J* = 245.6 Hz, C-4′), 137.0 (d, *J* = 2.4 Hz, C-1′), 130.3 (d, *J* = 8.1Hz, C-2′ and C-6′), 117.0 (d, *J* = 21.8 Hz, C-3′ and C-5′),103.9 (C-4a), 97.6 (C-6), 96.6 (C-8), 80.1 (C-2), 44.4 (C-3) ppm. Carbon atoms at positions 1′, 2′, 3′, 4′, 5′, and 6′ exhibited duplicated peaks, consistent with the influence of fluorine substitution on the aromatic B-ring[1].
HRMS (ESI$^-$): *m/z* calculated for $C_{15}H_{10}FO_4^-$, 273.05686 $[M-H]^-$; found 273.05570.
The annotated structure of 4′-fluoro-5,7-dihydroxyflavanone, including key 2D-NMR correlations, is shown in Figure S10. The $^1$H- and $^{13}$C-NMR spectra are shown in Figures S17 and S18, respectively.

### Supplementary Figures

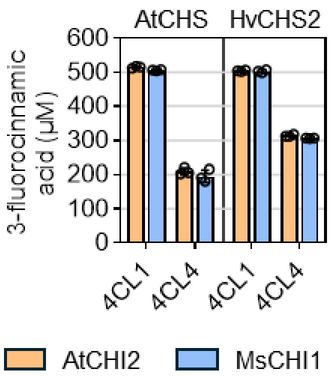


**Figure S1.** Remaining substrate titres in *Escherichia coli* strain SBC016072 carrying different pathway variants, quantified 24 h after supplementation with 3-fluorocinnamic acid at a final concentration of 0.5 mM. CHS and 4CL identities are indicated within the chart, while CHI identity is represented by bar colour (orange: AtCHI2; blue: MsCHI1). Data are presented as mean ± standard deviation ($n = 3$).

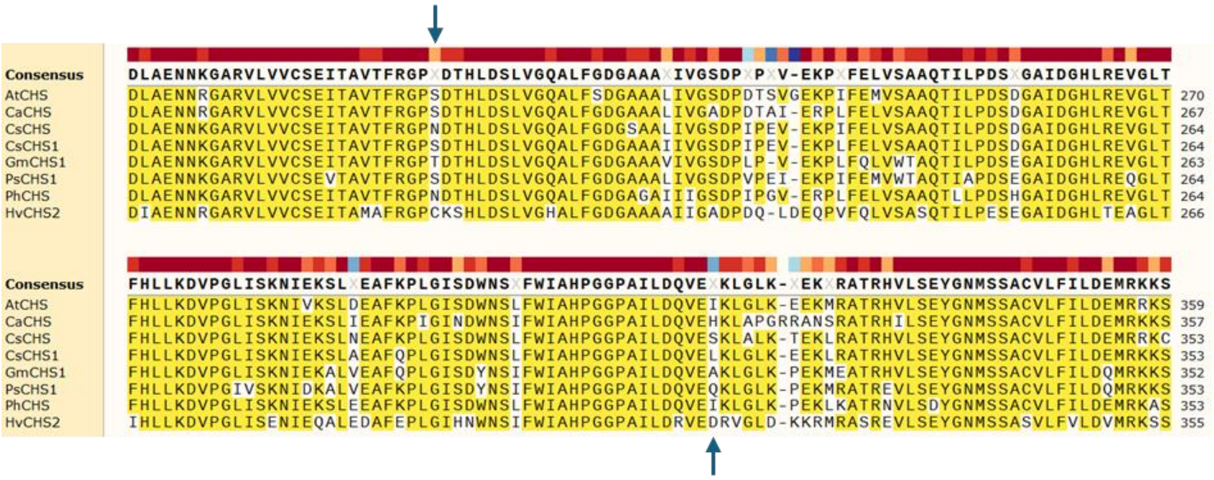


**Figure S2.** Multiple sequence alignment of chalcone synthases used for flavanone biosynthesis. Aligned protein sequences include CHS from *Arabidopsis thaliana* (AtCHS, Uniprot P13114), *Chrysosplenium americanum* (CaCHS, Uniprot O04220), *Cannabis sativa* (CsCHS, Uniprot Q8RVK9), CHS1 from *Camellia sinensis* (CsCHS1, P48386), CHS1 from *Glycine max* (GmCHS1, Uniprot P24826), CHS1 from *Pisum sativum* (PsCHS1, Uniprot Q01286), CHS from *Petunia hybrida* (PhCHS, Uniprot P08894), and CHS2 from *Hordeum vulgare* (HvCHS2, Uniprot Q96562). Sequences were aligned using Clustal Omega in SnapGene (version 4.3.11). The displayed region corresponds to HvCHS2 residues D178–S335. Sequence conservation is shown as coloured blocks, and residues with > 50% consensus are highlighted in yellow. Positions corresponding to HvCHS2 C204 and D317 are indicated.

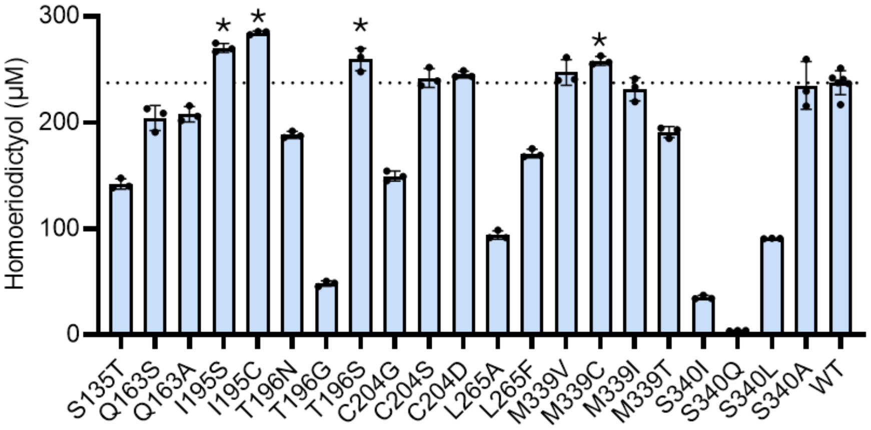


**Figure S3.** Homoeriodictyol titres in *E. coli* strain SBC016072 carrying plasmid SBC016421 with different HvCHS2 variants, measured 24 h after supplementation with 0.5 mM ferulic acid. Data are presented as mean ± standard deviation of three biological replicates, except for the parent enzyme (WT), which represents the mean ± standard deviation of six biological replicates. Statistical significance is indicated as * where $p < 0.05$ (unpaired two-tailed $t$ test).

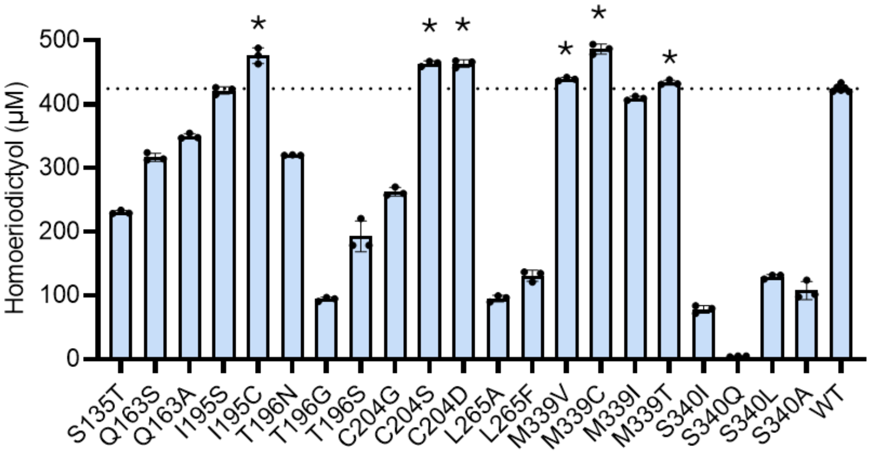


**Figure S4.** Homoeriodictyol titres in *E. coli* strain SBC016072 carrying plasmid SBC016421 with different HvCHS2 variants, measured 24 h after supplementation with 1.0 mM ferulic acid. Data are presented as mean ± standard deviation of three biological replicates, except for the parent enzyme (WT), which represents the mean ± standard deviation of six biological replicates. Statistical significance is indicated as * where $p < 0.05$ (unpaired two-tailed $t$ test).

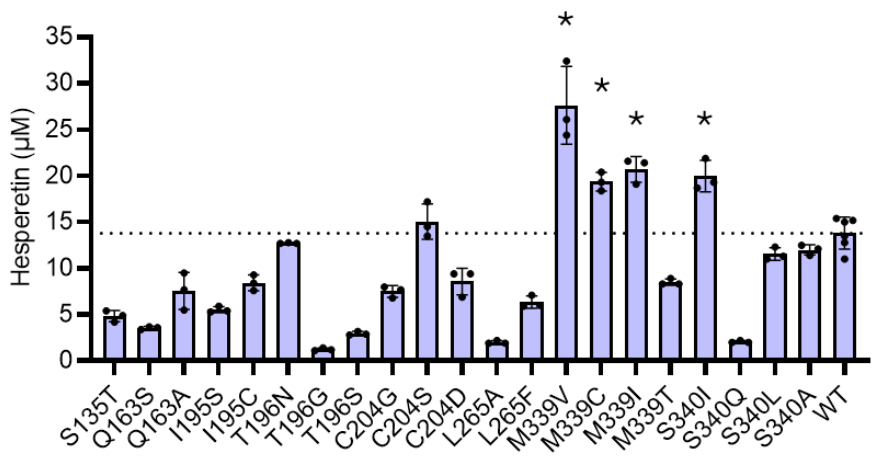


**Figure S5.** Hesperetin titres in *E. coli* strain SBC016072 carrying plasmid SBC016421 with different HvCHS2 variants, measured 24 h after supplementation with 0.5 mM isoferulic acid. Data are presented as mean ± standard deviation of three biological replicates, except for the parent enzyme (WT), which represents the mean ± standard deviation of six biological replicates. Statistical significance is indicated as * where $p < 0.05$ (unpaired two-tailed $t$ test).

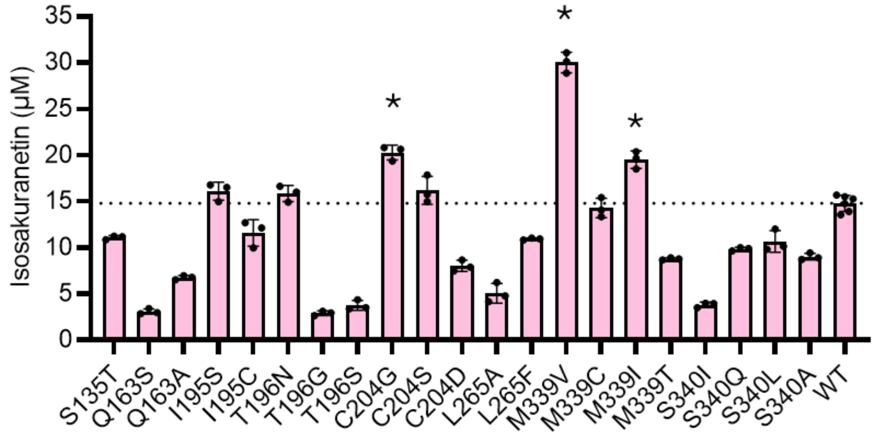


**Figure S6.** Isosakuranetin titres in *E. coli* strain SBC016072 carrying plasmid SBC016421 with different HvCHS2 variants, measured 24 h after supplementation with 0.5 mM 4-methoxycinnamic acid. Data are presented as mean ± standard deviation of three biological replicates, except for the parent enzyme (WT), which represents the mean ± standard deviation of six biological replicates. Statistical significance is indicated as * where $p < 0.05$ (unpaired two-tailed $t$ test).

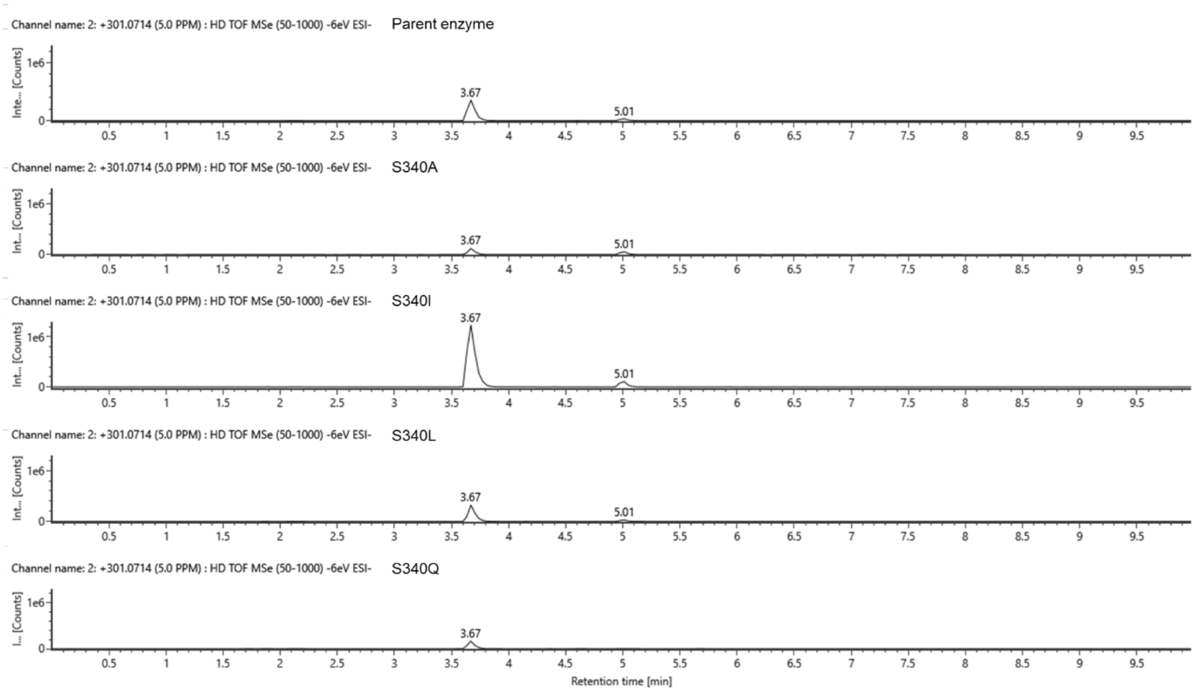


**Figure S7.** Extracted ion chromatogram (*m*/*z* 301.0714; $MS^E$ $ESI^-$, 6 eV) of culture extracts from the parent HvCHS2 strain and S340A, S340I, S340L, and S340Q variants. Peaks at retention times of 3.67 min and 5.01 min correspond to derailment product A and hesperetin, respectively.

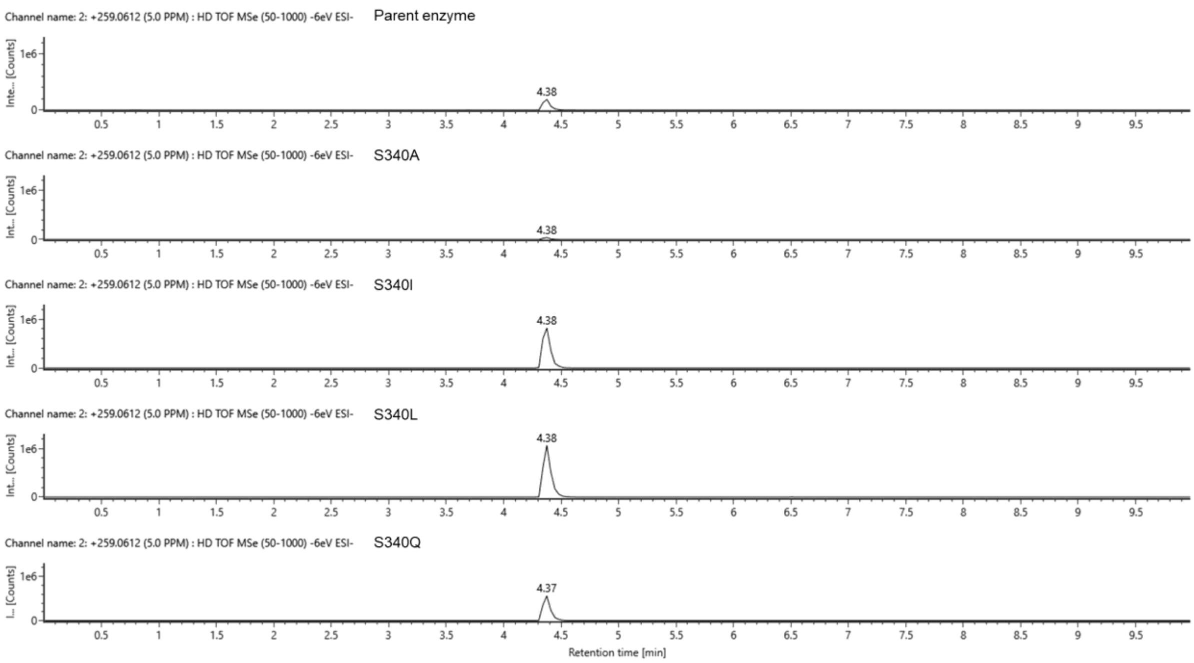


**Figure S8.** Extracted ion chromatogram (*m*/*z* 259.0612; $MS^E$ $ESI^-$, 6 eV) of culture extracts from the parent HvCHS2 strain and S340A, S340I, S340L, and S340Q variants. The peak at retention time of 4.38 min corresponds to derailment product B.

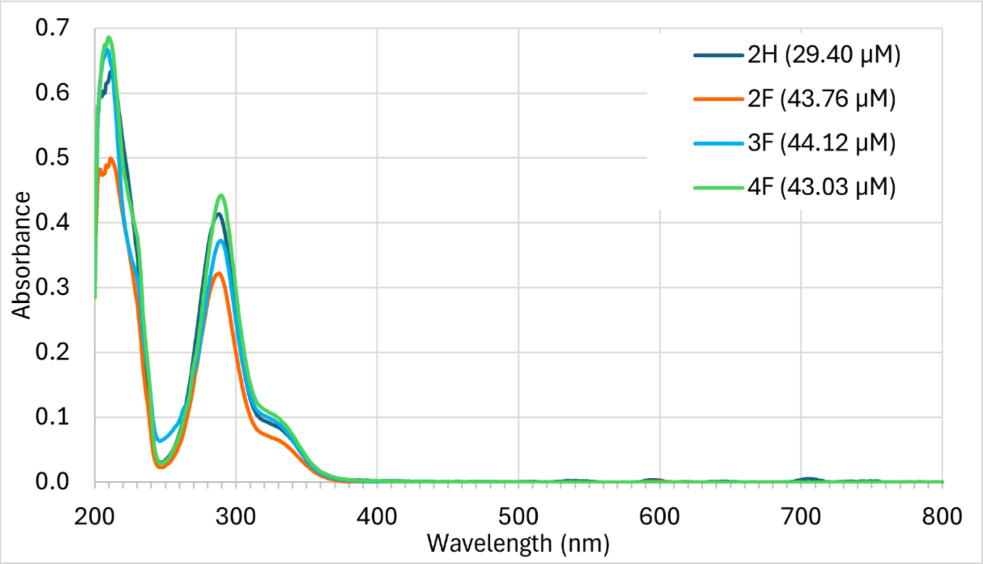


**Figure S9.** UV-Vis spectra of isolated flavanones. Spectra were recorded in 100% methanol at room temperature using a path length of 10 mm. Flavanones shown are 5,7,2′-trihydroxyflavanone (2H, dark blue spectrum), 2′-fluoro-5,7-dihydroxyflavanone (2F, orange spectrum), 3′-fluoro-5,7-dihydroxyflavanone (3F, light blue spectrum), and 4′-fluoro-5,7-dihydroxyflavanone (4F, green spectrum).

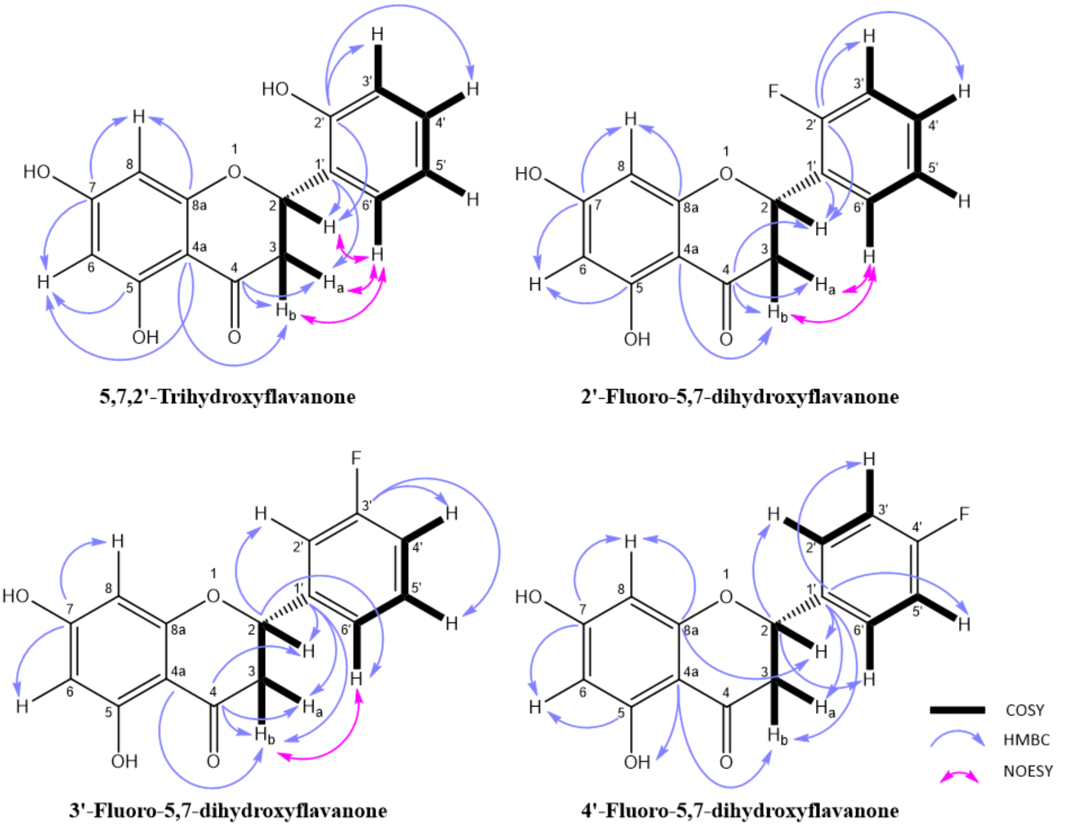


**Figure S10.** Annotated structures and key 2D-NMR correlations of isolated flavanones, including 5,7,2′-trihydroxyflavanone, 2′-fluoro-5,7-dihydroxyflavanone, 3′-fluoro-5,7-dihydroxyflavanone, and 4′-fluoro-5,7-dihydroxyflavanone.

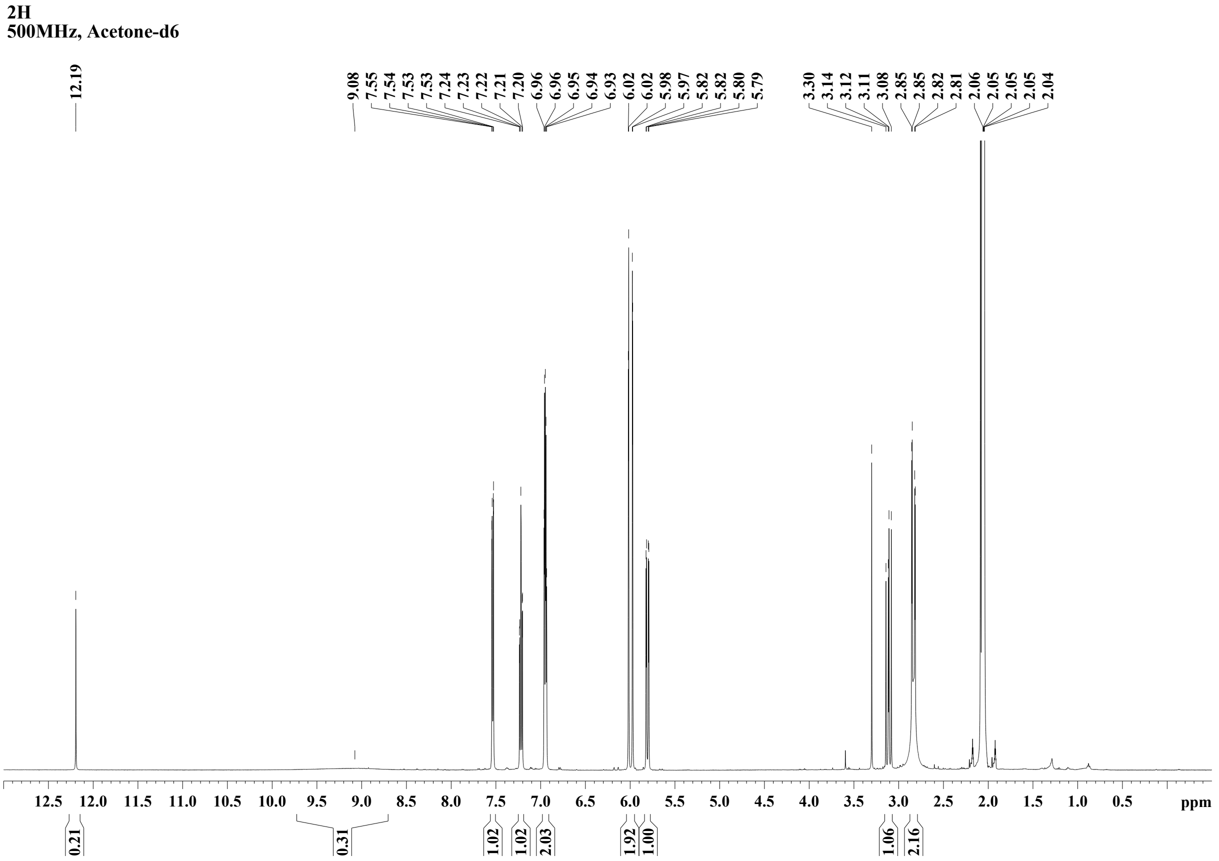


**Figure S11.** [1]H-NMR spectrum of 5,7,2′-trihydroxyflavanone.

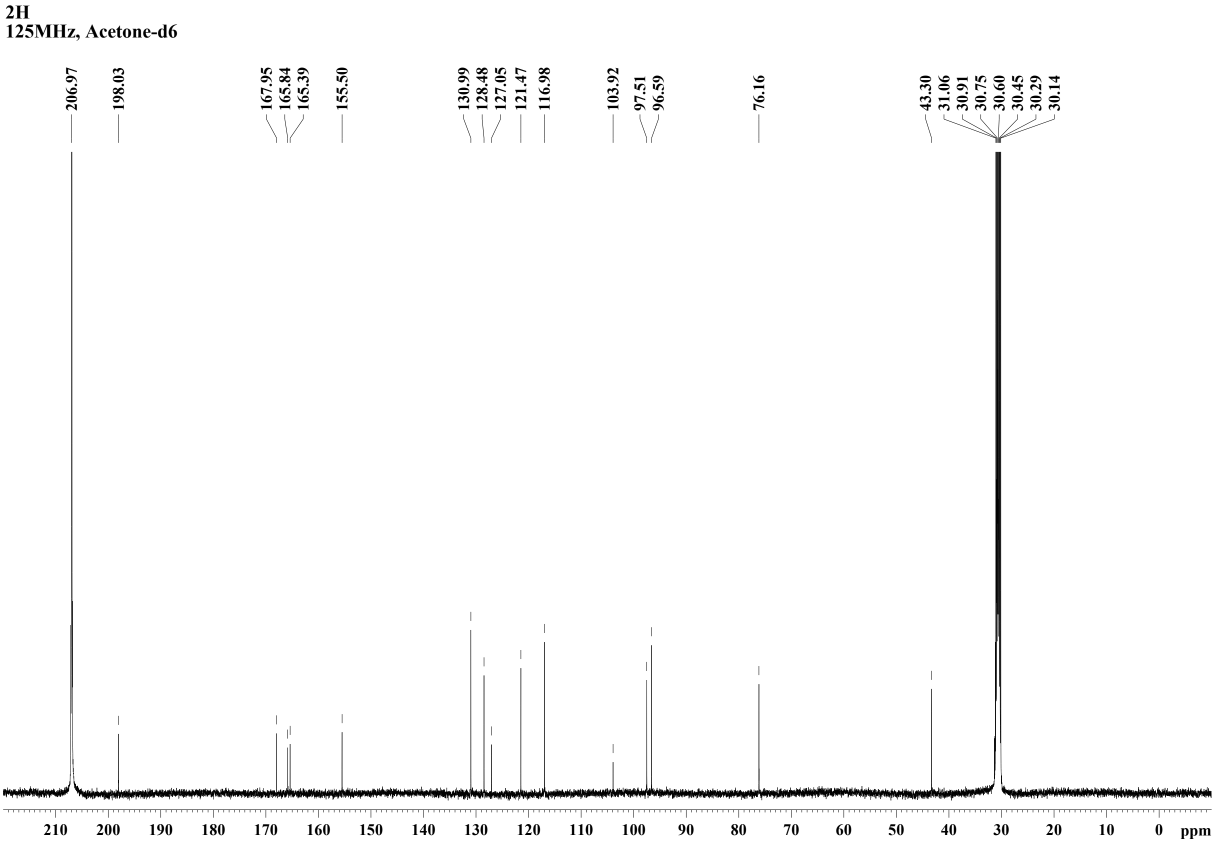


**Figure S12.** $^{13}$C-NMR spectrum of 5,7,2′-trihydroxyflavanone.

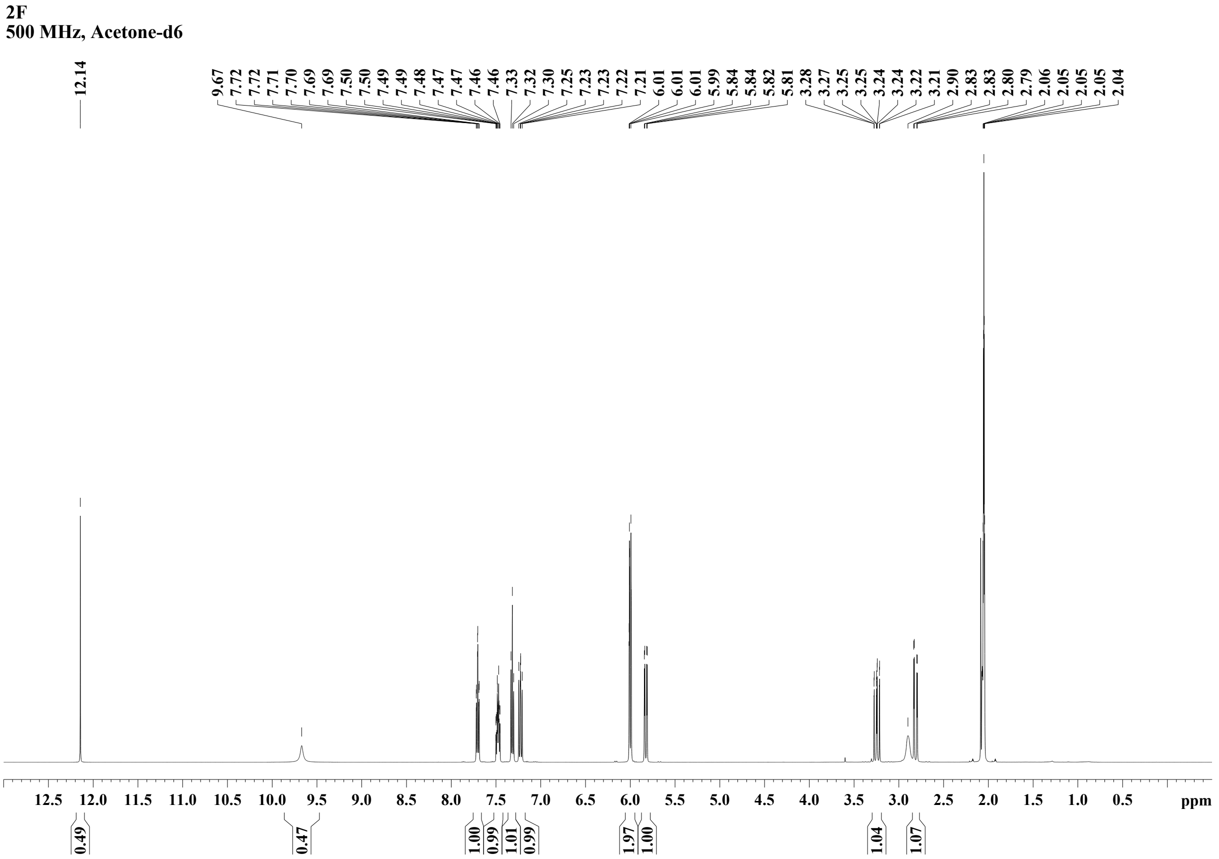


**Figure S13.** $^{1}$H-NMR spectrum of 2′-fluoro-5,7-dihydroxyflavanone.

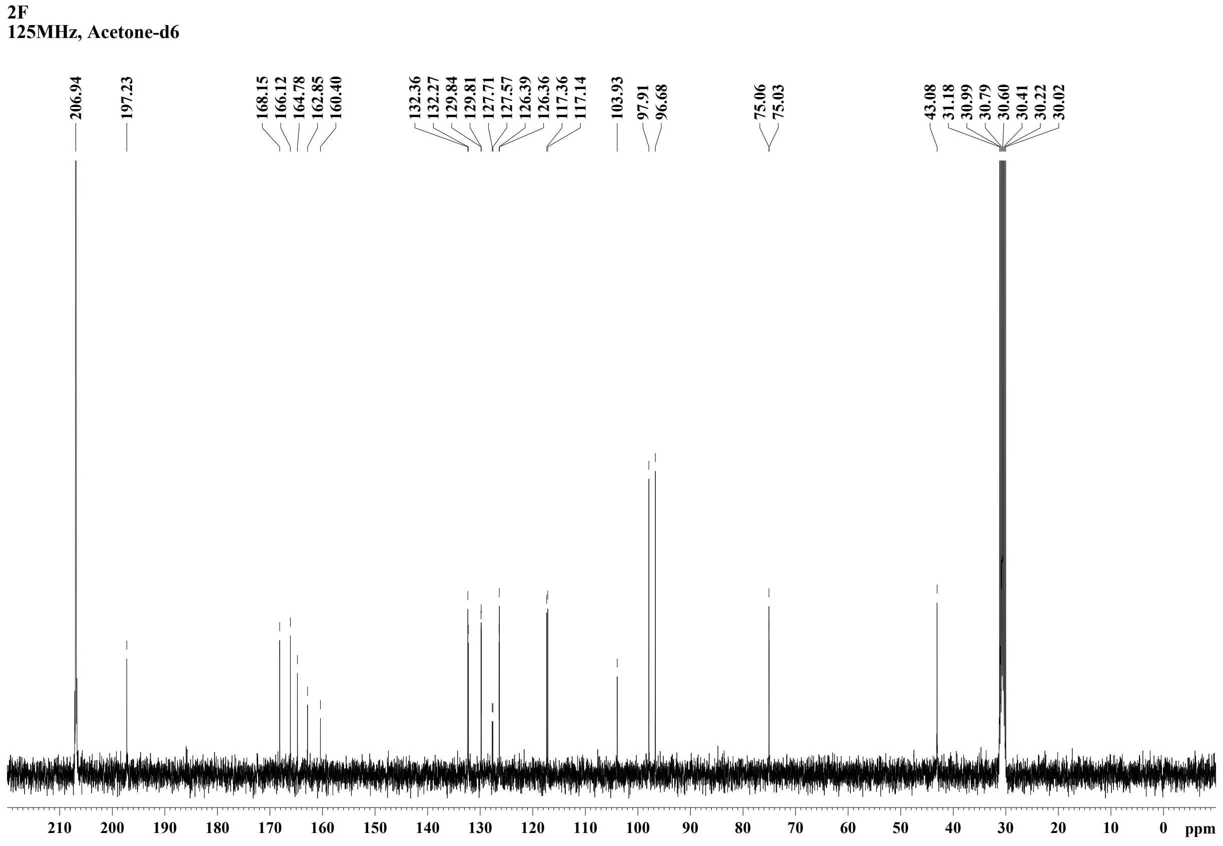


**Figure S14.** $^{13}$C-NMR spectrum of 2′-fluoro-5,7-dihydroxyflavanone.

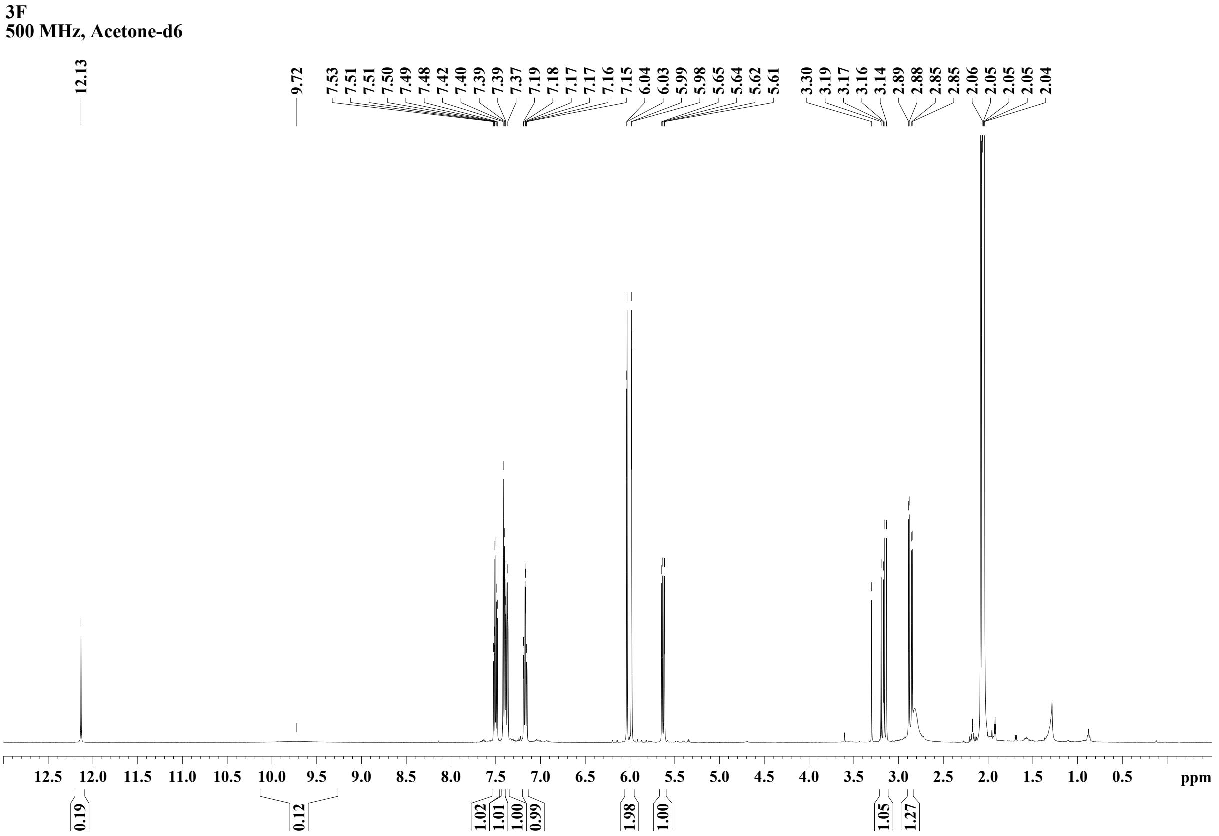


**Figure S15.** $^{1}$H-NMR spectrum of 3′-fluoro-5,7-dihydroxyflavanone.

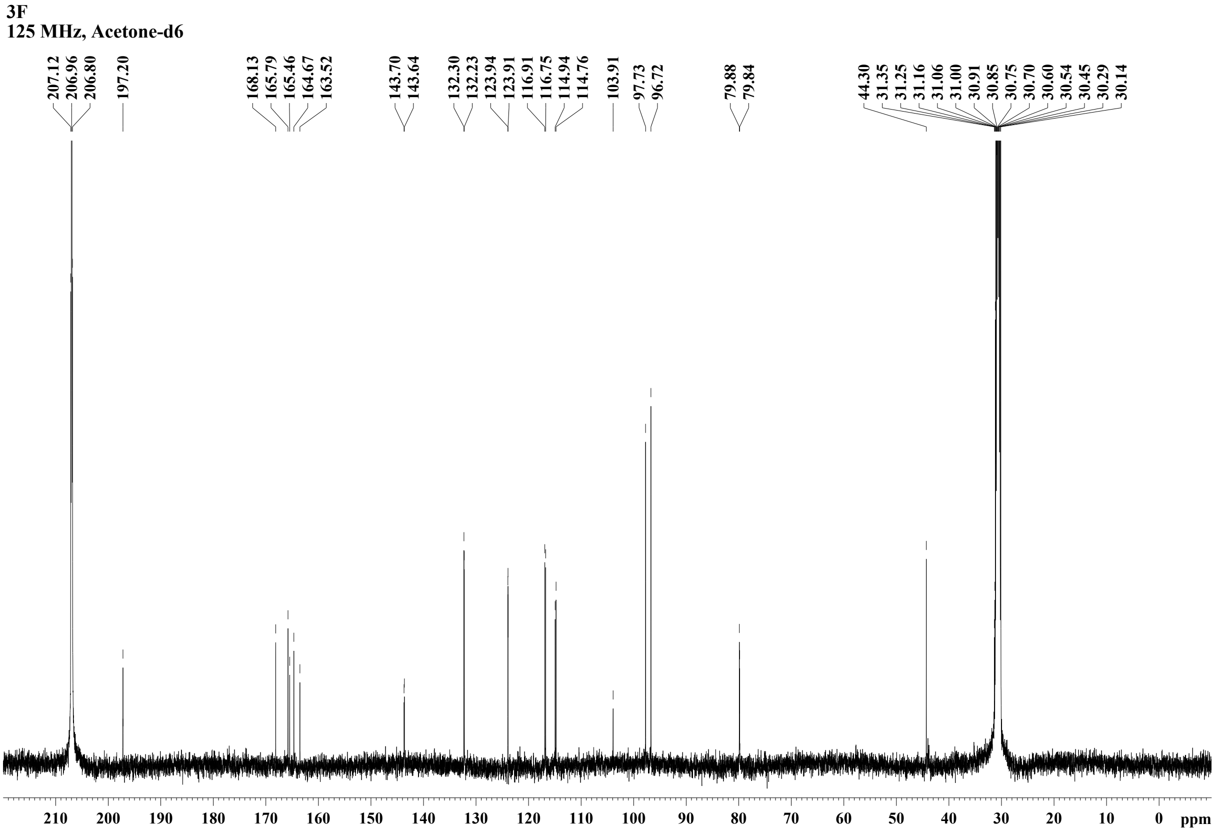


**Figure S16.** $^{13}$C-NMR spectrum of 3′-fluoro-5,7-dihydroxyflavanone.

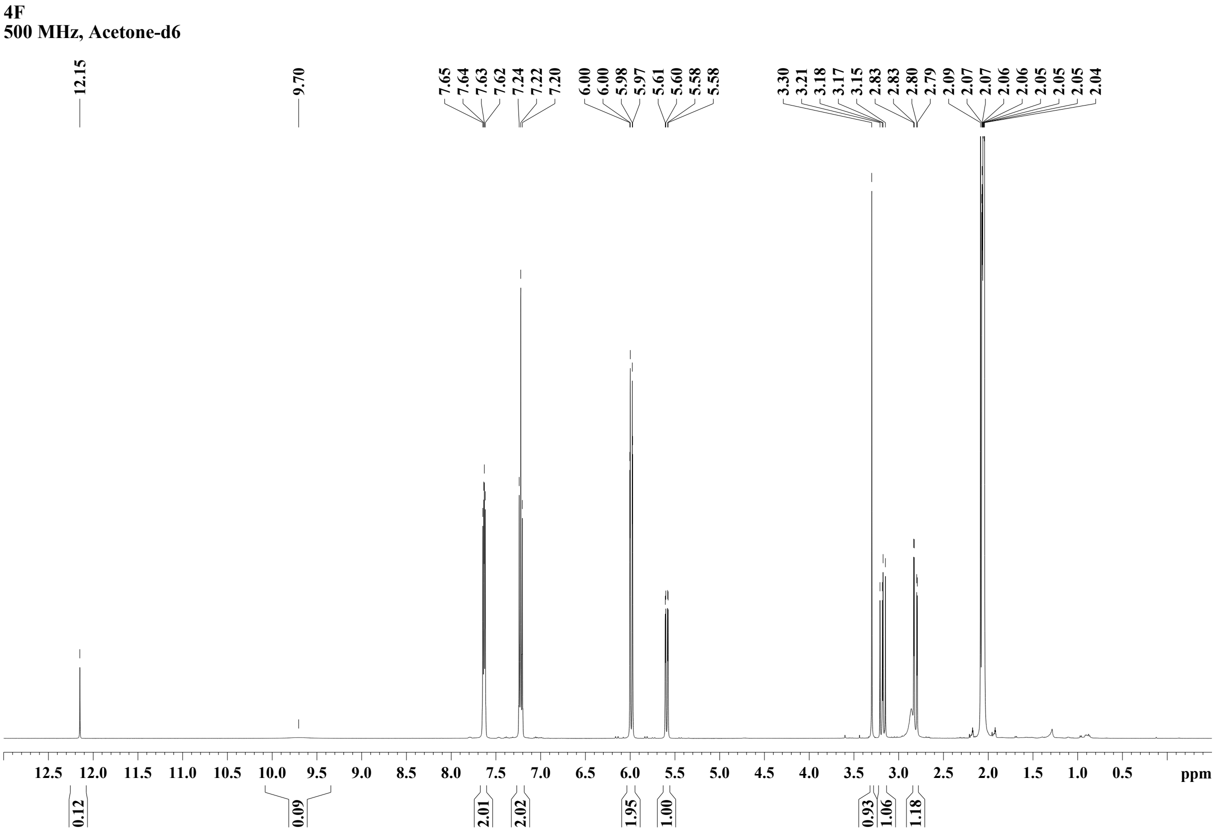


**Figure S17.** $^{1}$H-NMR spectrum of 4′-fluoro-5,7-dihydroxyflavanone.

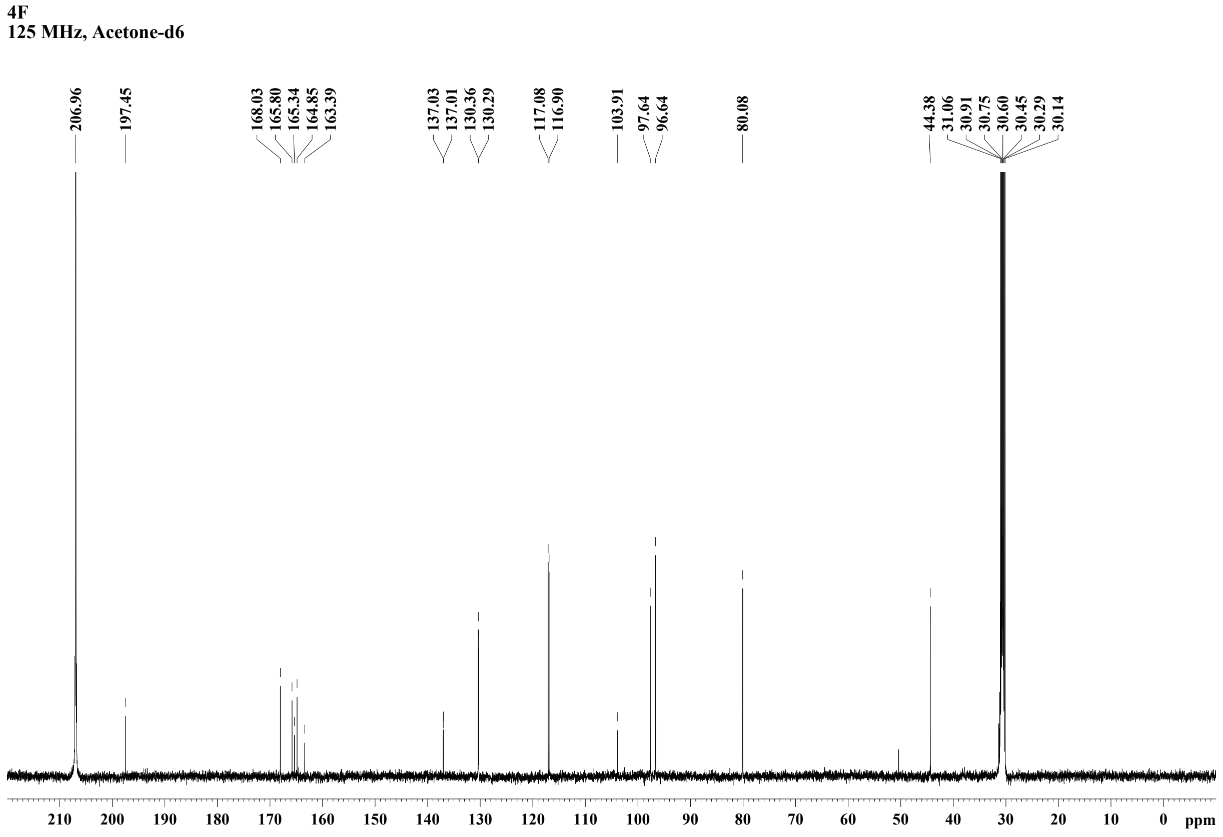


**Figure S18.** $^{13}$C-NMR spectrum of 4′-fluoro-5,7-dihydroxyflavanone.

#### Supplementary Tables

**Table S1.** Ring-substituted cinnamic acid and flavanone analytical standards used in this study.

| **Chemical** | **Manufacturer (catalogue number)** |
|---|---|
| Cinnamic acid | Sigma-Aldrich (C80857) |
| 2-Hydroxycinnamic acid | Sigma-Aldrich (H22809) |
| 3-Hydroxycinnamic acid | Sigma-Aldrich (H23007) |
| *p*-Coumaric acid | Sigma-Aldrich (C9008) |
| 2-Fluorocinnamic acid | Sigma-Aldrich (222712) |
| 3-Fluorocinnamic acid | Fluorochem (F001441) |
| 4-Fluorocinnamic acid | Sigma-Aldrich (222720) |
| 4-Methoxycinnamic acid | Sigma-Aldrich (M13807) |
| Caffeic acid | Sigma-Aldrich (C0625) |
| Isoferulic acid | Fluorochem (F791625) |
| Ferulic acid | Sigma-Aldrich (128708) |
| Pinocembrin | PhytoLab (PHL80061) |
| Naringenin | Sigma-Aldrich (N5893) |
| Isosakuranetin | PhytoLab (PHL82569) |
| Eriodictyol | Thermo Scientific Chemicals (465240010) |
| Hesperetin | PhytoLab (PHL89222) |
| Homoeriodictyol | Fluorochem (F830977) |

**Table S2.** Oligonucleotide primers used in this study. SSM – site saturation mutagenesis.

| **Primer name** | **Sequence (5′ to 3′)** | **Purpose** |
|---|---|---|
| EHfbrh_081 | GCTGGCAATTCCGACGTCTTAACAGATACCAGGTAAC | Plasmid assembly |
| EHfbrh_196 | CCGGTGCTTAAGCGGCCGCGTTTATCACAGTTAAATTGCTAACGCAGTCAGG | Plasmid assembly |
| EHfbrh_197 | TCACTCATTAGGCACCTCAGGTCGAGGTGGCCCG | Plasmid assembly |
| EHfbrh_215 | GCTTAAGCACCGGTGGAG | Plasmid assembly |
| EHfbrh_216 | GGTGCCTAATGAGTGAGCTAAC | Plasmid assembly |
| EHfbrh_217 | GCTCTGTTAATTAACGTCTTCGCTTCCCTACG | Plasmid assembly |
| EHfbrh_218 | GACGTTAATTAACAGAGCTAAAACAAACACGTAAATTACC | Plasmid assembly |
| EHfbrh_219 | TTCTCGGGTTTGGGCGCGCCTTACGGTGTCTGCGTCGC | Plasmid assembly |
| EHfbrh_220 | CCCAAACCCGAGAATCTCC | Plasmid assembly |
| EHfbrh_221 | TTTAGACCCTGCAGGGCCTGCGTTTACCGTGAG | Plasmid assembly |

| | | |
|---|---|---|
| EHfbrh_222 | CCTGCAGGGTCTAAATTACACGTCTCCTAGGGTT | Plasmid assembly |
| EHfbrh_223 | TCGGAATTGCCAGCTGGG | Plasmid assembly |
| EHfbrh_224 | TTCAATAGATACCGCCGTAGG | Plasmid assembly |
| EHfbrh_256 | TAAACGCAGGCCCTGCAG | SSM |
| EHfbrh_257 | GCCTGGAGATCCTTACTCGAG | SSM |
| EHfbrh_284 | ATTCTCGGGTTTGGGCGCGCCTTAATTGGCCACAACCAG ACC | Plasmid assembly |
| EHfbrh_288 | AACGCAGGCCCTGCAGGAGTAGAGATGTACAGTGTAGAA GTGC | Plasmid assembly |
| EHfbrh_289 | AGATCCTTACTCGAGTTTGGATCCTTACGCACTCGCCGC AGT | Plasmid assembly |
| EHfbrh_290 | CGCTTAAACGTTCAAGACTTCTTAAAAGATCTTTTGAAT TCTGAAATTG | Plasmid assembly |
| EHfbrh_291 | TGCCTTTAAGATTGGTAACTAACAAACAAGAAGTAACTT ATCGGCAGT | Plasmid assembly |
| EHfbrh_292 | TCTTGAACGTTTAAGCGTGCT | Plasmid assembly |
| EHfbrh_293 | TTAGTTACCAATCTTAAAGGCACCCT | Plasmid assembly |
| EHfbrh_294 | AATTTCGGAACACACAACTAAGACG | SSM (HvCHS2 T196) |
| EHfbrh_295 | TCTTAGTTGTGTGTTCCGAAATTNNKGCGATGGCTTTCC GCGGA | SSM (HvCHS2 T196) |
| EHfbrh_315 | CCACCTGGTTTTCTGTACCNNKTCAGGCGTCGATATGCC T | SSM (HvCHS2 T134) |
| EHfbrh_316 | GGTACAGAAAACCAGGTGGG | SSM (HvCHS2 T134) |
| EHfbrh_317 | ACCTGGTTTTCTGTACCACTNNKGGCGTCGATATGCCTG GC | SSM (HvCHS2 S135) |
| EHfbrh_318 | AGTGGTACAGAAAACCAGGTGG | SSM (HvCHS2 S135) |
| EHfbrh_319 | CACTTCAGGCGTCGATNNKCCTGGCGCCGATTATCAG | SSM (HvCHS2 M139) |
| EHfbrh_320 | ATCGACGCCTGAAGTGGT | SSM (HvCHS2 M139) |
| EHfbrh_321 | GTGAAGCGTCTGATGATGTACNNKCAGGGGTGCTTTGGG GGC | SSM (HvCHS2 Q163) |
| EHfbrh_322 | GTACATCATCAGACGCTTCACTG | SSM (HvCHS2 Q163) |
| EHfbrh_323 | GCGTCTTAGTTGTGTGTTCCNNKATTACGGCGATGGCTT TCC | SSM (HvCHS2 E194) |
| EHfbrh_324 | GGAACACACAACTAAGACGCG | SSM (HvCHS2 E194) |
| EHfbrh_325 | GTCTTAGTTGTGTGTTCCGAANNKACGGCGATGGCTTTC CGC | SSM (HvCHS2 I195) |
| EHfbrh_326 | TTCGGAACACACAACTAAGACG | SSM (HvCHS2 I195) |
| EHfbrh_327 | CTTTCCGCGGACCGNNKAAGTCTCATCTAGATAGCCTGG TC | SSM (HvCHS2 C204) |
| EHfbrh_328 | CGGTCCGCGGAAAGCCAT | SSM (HvCHS2 C204) |
| EHfbrh_329 | CTGGTCGGTCATGCGNNKTTCGGGGATGGTGCCGCG | SSM (HvCHS2 L216) |

| | | |
|---|---|---|
| EHfbrh_330 | CGCATGACCGACCAGGCT | SSM (HvCHS2 L216) |
| EHfbrh_331 | TCATGCGTTGTTCGGGNNKGGTGCCGCGGCGGCTATT | SSM (HvCHS2 D219) |
| EHfbrh_332 | CCCGAACAACGCATGACC | SSM (HvCHS2 D219) |
| EHfbrh_333 | GAAAGCGAAGGGGCTNNKGATGGTCATTTGACTGAGGCG | SSM (HvCHS2 I256) |
| EHfbrh_334 | AGCCCCTTCGCTTTCCGG | SSM (HvCHS2 I256) |
| EHfbrh_337 | CGGGACTGACCATACATNNKCTGAAAGATGTGCCTGGC | SSM (HvCHS2 L269) |
| EHfbrh_338 | ATGTATGGTCAGTCCCGC | SSM (HvCHS2 L269) |
| EHfbrh_339 | ACTGACCATACATCTACTGAAAGATNNKCCTGGCCTCAT TTCCGAAA | SSM (HvCHS2 V273) |
| EHfbrh_340 | ATCTTTCAGTAGATGTATGGTCAGTCC | SSM (HvCHS2 V273) |
| EHfbrh_341 | ATCCTGGATCGTGTGGAANNKCGCGTAGGCTTGGATAAG AAA | SSM (HvCHS2 D317) |
| EHfbrh_342 | TTCCACACGATCCAGGATAGC | SSM (HvCHS2 D317) |
| EHfbrh_343 | CTCAGCGAGTATGGGAACNNKAGCTCAGCAAGCGTATTA TTTGTTC | SSM (HvCHS2 M339) |
| EHfbrh_344 | GTTCCCATACTCGCTGAGC | SSM (HvCHS2 M339) |
| EHfbrh_345 | CAGCGAGTATGGGAACATGNNKTCAGCAAGCGTATTATT TGTTCTGG | SSM (HvCHS2 S340) |
| EHfbrh_346 | CATGTTCCCATACTCGCTGAG | SSM (HvCHS2 S340) |
| EHfbrh_377 | AATCTCACTGCACACAACTAATACG | SSM (AtCHS T199) |
| EHfbrh_378 | GTATTAGTTGTGTGCAGTGAGATTNNKGCCGTTACCTTC CGCGGC | SSM (AtCHS T199) |
| EHfbrh_379 | ATTGCCATACTCGGACAGC | SSM (AtCHS M343) |
| EHfbrh_380 | CTGTCCGAGTATGGCAATNNKTCGAGTGCATGCGTGCTG | SSM (AtCHS M343) |
| EHfbrh_394 | CTCACTGCACACAACTAATACGC | SSM (AtCHS I198) |
| EHfbrh_395 | CGTATTAGTTGTGTGCAGTGAGNNKACTGCCGTTACCTT CCGC | SSM (AtCHS I198) |

**Table S3.** Sequences of the synthesised gene parts. Gene coding sequences are shown in uppercase letters. Restriction enzyme recognition sites are italicised.

| Gene part | Sequence |
|---|---|
| FdeR | *gacgtc*TTAACAGATACCAGGTAACGCAGCGTCCATCTCCTGAGCACTCTCTAAGAAG ACGCGGCGTAACCACTGAATTCCGGGATCATTGGAGCGGTAGCGATGCCATTGCATCA TTTGGCGCATTTCTCCAAGTGACAAAGGACTCTCCTTGATGACTACCGGCCATTGCGG TGCCAACAGTTGGGCTAAGCGAGCGTGCACCGTGGCGATGCGGTCCGTACCTTGGACC AGAGCAAGAGCAGATGCGAATGAAAATGACGTAACCTCCACGCGGCGAGCAAATCCCA ACTTACGAGCCATCCACGCTTCGACCGAGGACGCGTTGGCGCCTGGTGGCACCATCAC CACATGACCGCTAGCCATGTAGCGCTCCAGAGTAAGCTCCCCCTGTGCCAATGCGGAA TCACGCCATACTACGCAAACATGACGTTCACGGAACACCTCCTCGGCGGGGTGATCTG GGGTACAAAACTCCTGCGGTAAAACCAATAAGTCAACTTCCGCGCGGTCCAAGGAGCG CGTTGGGTCCTGAACCTGTGGCATCAGAGCAAAGCGAATGTGCTTGCCCTCAGCATGT |

| | |
|---|---|
| | GCACGTGCAAGCACGCGGGGAATCAGGACGCTCAATGTAAAGTCCGAGACGCTAATAC<br>GAAACTCACGGGTCGACTCTGCAGGAACAAAAGCAGGCAAGGCAGCGATAGAGCCGTC<br>GATGCGACGTAAAACATCATGCACCGCGTCTTTTAATACTTCTGCACGTGGAGTAGGT<br>TCCATGCGGCGCCCGACCTGAATAAGCAGTTCATCGTCGAAATACTCACGAAGACGTG<br>CTAAGGCATTGGACATCGCGGATTGAGACAAGTGAATTTTTTCCGCAGCACGGCTGAT<br>GCTCATCTCAGTAAGAAGCGCGTCCAGAGCCACCAGTAAATTCAGGTCCAGCTTATTA<br>AAGCGCATggcgctggtctccgttgttgtgcttgttcttgccgaccctcggatagacg<br>acggatggggtggtcaatgtattgatgccgtccatatcatgaatcaaaacaatccatt<br>tgatcaatatcaagctcactcttaagcttcactcatccgctgcatggccccaccagaa<br>agggctggcgcggcaagccggcggcgcactcgcactggatgcgccgctgttgagcctg<br>gccatgacaacgcgccgatagcggccacaccccgccaggcagggtaggagacaaggag<br>aca*catatg* |
| Gm4CL1 | ttcaatagataccgccgtagggaagcgaagacgttaattaaaactggccgtattgtta<br>tctgcgtgggtacccacaccaatactttccaggaggtaattacATGGCGCCTTCGCCG<br>CAAGAAATCATTTTTCGCTCGCCACTTCCGGATATTCCGATCCCGACACATTTACCGC<br>TCTATTCCTACTGCTTTCAGAATCTTTCACAGTTCCATGATCGCCCTTGTCTTATTGA<br>CGGAGACACCGGCGAAACGCTGACATATGCAGACGTGGACCTGGCGGCCCGCCGTATT<br>GCGAGCGGCCTGCATAAGATTGGTATCAGACAAGGTGATGTTATTATGCTAGTCCTCC<br>GGAACTGTCCTCAATTCGCCTTAGCCTTTCTGGGAGCGACACACCGTGGGGCGGTAGT<br>GACCACTGCAAATCCCTTTTATACGCCCGCGGAGTTAGCCAAACAGGCGACAGCAACC<br>AAAACCCGCCTGGTGATTACCCAAAGCGCCTACGTGGAAAAGATTAAAAGCTTTGCTG<br>ACAGCAGTAGCGACGTAATGGTTATGTGCATCGACGATGACTTTTCCTATGAAAACGA<br>TGGTGTTCTTCACTTCAGCACACTAAGTAACGCCGATGAGACGGAAGCCCCAGCGGTG<br>AAGATTAACCCTGATGAGCTGGTAGCCCTGCCATTTAGTTCAGGGACGTCTGGCCTAC<br>CAAAAGGCGTGATGCTTAGCCACAAAAACCTGGTGACCACAATTGCCCAACTGGTTGA<br>TGGCGAGAACCCGCACCAGTATACACATTCTGAAGATGTGCTATTGTGCGTGCTGCCT<br>ATGTTTCATATCTACGCCCTGAACTCCATCCTGTTGTGCGGTATCCGTTCAGGCGCCG<br>CGGTGCTCATTCTGCAGAAATTCGAAATTACCACTCTGCTGGAACTAATCGAGAAATA<br>CAAGGTGACGGTGGCCAGTTTCGTTCCTCCGATTGTCTTAGCCTTGGTTAAATCAGGC<br>GAGACACACCGATATGATCTGAGTTCCATTCGCGCAGTAGTTACGGGCGCAGCCCCGT<br>TAGGCGGAGAACTGCAAGAGGCCGTCAAAGCACGCCTGCCTCACGCAACCTTTGGCCA<br>GGGCTATGGTATGACTGAAGCAGGCCCGCTGGCGATTAGCATGGCCTTTGCCAAAGTA<br>CCGTCGAAAATCAAGCCAGGGGCGTGCGGTACCGTAGTGCGCAATGCGGAGATGAAAA<br>TTGTAGACACCGAAACCGGCGATAGTCTACCGCGAAATAAACACGGCGAAATTTGCAT<br>TATTGGCACTAAAGTGATGAAAGGGTATCTCAACGACCCGGAAGCCACTGAACGCACC<br>GTCGATAAAGAAGGCTGGTTACACACCGGAGATATTGGTTTTATTGATGATGACGATG<br>AATTATTCATTGTTGATCGCTTAAAAGAACTAATCAAATACAAAGGTTTTCAAGTGGC<br>CCCGGCCGAACTGGAAGCCCTGCTGATTGCACACCCAAATATTTCGGATGCAGCCGTT<br>GTGGGGATGAAAGATGAAGCTGCTGGTGAAATACCCGTGGCATTTGTCGTGCGAAGTA<br>ATGGCAGCGAAATTGCCGAGGATGAGATCAAAAAGTATATTAGCCAGCAAGTGGTGTT<br>TTACAAACGTATCTGTCGCGTGTTTTTCACCGACAGTATTCCTAAGGCGCCGTCTGGC<br>AAGATTCTGCGTAAAGTTCTCACGGCGCGCCTGAACGAAGGTCTGGTTGTGGCCAATT<br>AAggcgcgcccaaacccgagaat |
| HvCHS2 | agtagagatgtacagtgtagaagtgcatcagtggtagacacgggggaggtagcacaat<br>atATGGCAGCAGTTCGACTTAAGGAAGTACGCATGGCGCAGCGCGCCGAAGGCCTAGC<br>CACTGTCCTTGCCATTGGCACCGCGGTGCCAGCAAATTGTGTCTATCAGGCGACCTAT<br>CCGGATTATTATTTTCGCGTAACCAAAAGCGAGCACTTGGCGGACTTGAAGGAAAAAT<br>TCCAGCGAATGTGTGACAAATCTATGATCCGTAAACGTCACATGCATCTCACTGAGGA<br>AATTCTGATCAAGAATCCGAAGATTTGTGCGCATATGGAAACCAGTCTGGATGCTCGA<br>CACGCGATTGCTCTGGTGGAGGTGCCTAAGCTGGGACAAGGCGCGGCGGAAAAGGCGA<br>TTAAAGAGTGGGGCCAGCCGCTGTCTAAAATCACCCACCTGGTTTTCTGTACCACTTC<br>AGGCGTCGATATGCCTGGCGCCGATTATCAGTTAACCAAGTTACTGGGTCTGAGCCCC<br>ACAGTGAAGCGTCTGATGATGTACCAACAGGGGTGCTTTGGGGGCGCGACAGTTCTGC |

| | |
|---|---|
| | GGCTAGCAAAGGATATCGCGGAAAATAACCGTGGTGCGCGCGTCTTAGTTGTGTGTTC<br>CGAAATTACGGCGATGGCTTTCCGCGGACCGTGCAAGTCTCATCTAGATAGCCTGGTC<br>GGTCATGCGTTGTTCGGGGATGGTGCCGCGGCGGCTATTATCGGTGCCGATCCAGACC<br>CTTTAGTGGAACAACCTGTGTTTCAGTTAGTCAGCGCGTCACAAACTATATTACCGGA<br>AAGCGAAGGGGCTATCGATGGTCATTTGACTGAGGCGGGACTGACCATACATCTACTG<br>AAAGATGTGCCTGGCCTCATTTCCGAAAATATTGAGCAGGCGCTTGAGGATGCCTTTG<br>AGCCGCTCGGCATTCATAACTGGAACAGCATCTTTTGGATTGCGCATCCGGGTGGCCC<br>AGCTATCCTGGATCGTGTGGAAGACCGCGTAGGCTTGGATAAGAAACGCATGCGAGCG<br>TCACGCGAAGTGCTCAGCGAGTATGGGAACATGAGCTCAGCAAGCGTATTATTTGTTC<br>TGGACGTTATGCGCAAGTCAAGCGCCAAAGACGGGCTGGCCACAACCGGTGAAGGCAA<br>AGATTGGGGTGTGCTGTTCGGGTTTGGGCCTGGGTTAACAGTTGAAACCTTAGTGTTA<br>CATTCAGTGCCGGTACCGGTACCGACTGCGGCGAGTGCGTAA |
| MsCHI | tcttgaacgtttaagcgtgctgcggaagacacaagtttttcgttaatctaggaggttg<br>atATGGCCGCCAGCATCACTGCAATTACAGTAGAAAATCTCGAATACCCGGCTGTTGT<br>GACCTCTCCCGTCACCGGAAAATCATACTTCCTGGGTGGCGCTGGCGAACGCGGGCTC<br>ACCATCGAAGGCAATTTCATTAAGTTTACAGCGATTGGAGTTTATCTTGAAGATATTG<br>CTGTAGCTTCTCTGGCCGCCAAATGGAAGGGGAAGAGCTCAGAAGAACTGCTGGAAAC<br>TCTCGATTTTTACCGCGACATTATATCTGGCCCCTTCGAGAAACTGATTCGAGGATCG<br>AAAATACGTGAACTTAGCGGTCCAGAATACTCGCGCAAAGTGATGGAAAACTGTGTTG<br>CGCACTTGAAATCCGTAGGGACCTATGGCGATGCGGAAGCCGAAGCGATGCAGAAGTT<br>CGCCGAAGCCTTTAAGCCGGTGAATTTTCCGCCTGGGGCTTCGGTCTTTTACCGCCAA<br>AGCCCGGATGGCATTCTCGGACTGTCTTTTTCGCCCGATACCTCGATCCCGGAGAAAG<br>AGGCCGCCCTGATTGAAAACAAGGCGGTGAGCAGTGCGGTACTTGAAACCATGATCGG<br>CGAGCATGCGGTTAGTCCCGATCTGAAACGGTGCCTGGCCGCCCGCCTGCCGGCTCTC<br>CTGAACGAGGGTGCCTTTAAGATTGGTAACTAA |

**Table S4.** Plasmids used and generated in this study.

| Plasmid ID | Characteristic | Reference or source |
|---|---|---|
| pBbE1c-rfp | Chl$^{R}$, ColE1, P$_{trc}$-*rfp*-T$_{rrnB1}$ | [2] |
| pCAT207 | Tet$^{R}$, pBBR1, P$_{j5}$-*rfp*-T$_{rrnB1}$ | Addgene plasmid # 134884[3] |
| SBC006845 | Kan$^{R}$, ColE1, P$_{lacUV5}$-AtCHI-T$_{B1006}$-P$_{trc}$-Gm4CL4-T$_{B1006}$-P$_{trc}$-AtCHS-T$_{rrnB1}$ | [4] |
| SBC016239 | Chl$^{R}$, ColE1, P$_{fdeR}$-*fdeR*, P$_{fdeA}$-*rfp*-T$_{rrnB1}$ | This work |
| SBC016302 | Kan$^{R}$, ColE1, P$_{fdeR}$-*fdeR*, P$_{fdeA}$-*rfp*, P$_{lacUV5}$-AtCHI2-T$_{B1006}$-P$_{trc}$-Gm4CL4-T$_{B1006}$-P$_{trc}$-AtCHS-T$_{rrnB1}$ | This work |
| SBC016397 | Kan$^{R}$, ColE1, P$_{fdeR}$-*fdeR*, P$_{fdeA}$-*rfp*, P$_{lacUV5}$-AtCHI2-T$_{B1006}$-P$_{trc}$-Gm4CL1-T$_{B1006}$-P$_{trc}$-AtCHS-T$_{rrnB1}$ | This work |
| SBC016420 | Kan$^{R}$, ColE1, P$_{fdeR}$-*fdeR*, P$_{fdeA}$-*rfp*, P$_{lacUV5}$-AtCHI2-T$_{B1006}$-P$_{trc}$-Gm4CL1-T$_{B1006}$-P$_{trc}$-HvCHS2-T$_{rrnB1}$ | This work |
| SBC016421 | Kan$^{R}$, ColE1, P$_{fdeR}$-*fdeR*, P$_{fdeA}$-*rfp*, P$_{lacUV5}$-MsCHI1-T$_{B1006}$-P$_{trc}$-Gm4CL1-T$_{B1006}$-P$_{trc}$-HvCHS2-T$_{rrnB1}$ | This work |
| SBC016434 | Kan$^{R}$, ColE1, P$_{fdeR}$-*fdeR*, P$_{fdeA}$-*rfp*, P$_{lacUV5}$-MsCHI1-T$_{B1006}$-P$_{trc}$-Gm4CL1-T$_{B1006}$-P$_{trc}$-AtCHS-T$_{rrnB1}$ | This work |
| SBC016441 | Kan$^{R}$, ColE1, P$_{fdeR}$-*fdeR*, P$_{fdeA}$-*rfp*, P$_{lacUV5}$-AtCHI2-T$_{B1006}$-P$_{trc}$-Gm4CL4-T$_{B1006}$-P$_{trc}$-HvCHS2-T$_{rrnB1}$ | This work |

| | | |
|---|---|---|
| SBC016442 | Kan$^R$, ColE1, $P_{fdeR}$-*fdeR*, $P_{fdeA}$-*rfp*, $P_{lacUV5}$-MsCHI1-$T_{B1006}$-$P_{trc}$-Gm4CL4-$T_{B1006}$-$P_{trc}$-HvCHS2-$T_{rrnB1}$ | This work |
| SBC016443 | Kan$^R$, ColE1, $P_{fdeR}$-*fdeR*, $P_{fdeA}$-*rfp*, $P_{lacUV5}$-MsCHI1-$T_{B1006}$-$P_{trc}$-Gm4CL4-$T_{B1006}$-$P_{trc}$-AtCHS-$T_{rrnB1}$ | This work |

**Table S5.** MRM transitions and MS/MS operating parameters for LC-MS/MS analysis of target flavanones.

| Compound | Ionisation mode | Parent *m*/*z* | Daughter *m*/*z* | Cone voltage [V] | Collision energy [eV] |
|---|---|---|---|---|---|
| Hesperetin | Negative | 301.11 | 164.10 | 4 | 24 |
| Homoeriodictyol | Negative | 301.18 | 151.04 | 62 | 16 |
| Isosakuranetin | Negative | 285.12 | 164.04 | 8 | 24 |

**Table S6.** Contributions of individual enzymes and their interactions to flavanone formation. Product titres were analysed using three-way ANOVA with 4CL, CHS, and CHI treated as categorical factors. Values represent the percentage of total variation explained by each factor and interaction term, with corresponding *p*-values shown in parentheses.

| | **% of total variation (*p*-value)** | | | | | | |
|---|---|---|---|---|---|---|---|
| **Compound** | **4CL** | **CHS** | **CHI** | **4CL × CHS** | **4CL ×CHI** | **CHS × CHI** | **4CL × CHS × CHI** |
| Pinocembrin | 33.03 (<0.0001) | 45.55 (<0.0001) | 0.99 (<0.0001) | 19.59 (<0.0001) | 0.15 (0.0001) | 0.53 (<0.0001) | 0.07 (0.0031) |
| 5,7,2′-Trihydroxy-flavanone | 1.04 (0.0007) | 88.46 (<0.0001) | 4.86 (<0.0001) | 1.04 (0.0007) | 0.29 (0.0418) | 3.04 (<0.0001) | 0.32 (0.0331) |
| Naringenin | 60.61 (<0.0001) | 27.42 (<0.0001) | 0.20 (0.0485) | 6.10 (<0.0001) | 4.24 (<0.0001) | 0.38 (0.0096) | 0.33 (0.0150) |
| 2′-fluoro-5,7-dihydroxy-flavanone | 37.91 (<0.0001) | 35.06 (<0.0001) | 1.49 (<0.0001) | 23.35 (<0.0001) | 0.86 (<0.0001) | 0.70 (<0.0001) | 0.45 (<0.0001) |
| 3′-fluoro-5,7-dihydroxy-flavanone | 77.31 (<0.0001) | 7.33 (<0.0001) | 2.25 (0.0008) | 7.33 (<0.0001) | 2.25 (0.0008) | 0.70 (0.035) | 0.70 (0.035) |
| 4′-fluoro-5,7-dihydroxy-flavanone | 88.11 (<0.0001) | 6.78 (<0.0001) | 0.34 (<0.0001) | 4.24 (<0.0001) | 0.03 (0.0955) | 0.18 (0.0008) | 0.14 (0.0027) |
| Isosakura-netin | 0.84 (0.0147) | 94.77 (<0.0001) | 0.86 (0.0137) | 0.84 0.0147) | 0.005 (0.8331) | 0.86 (0.0137) | 0.005 (0.8331) |

| | | | | | | | |
|---|---|---|---|---|---|---|---|
| Eriodictyol | 31.97 (<0.0001) | 0.18 (0.5640) | 19.90 (<0.0001) | 13.23 (0.0001) | 9.16 (0.0006) | 1.82 (0.0760) | 15.65 (<0.0001) |
| Hesperetin | 0.86 (0.2385) | 85.09 (<0.0001) | 1.97 (0.0826) | 0.86 (0.2385) | 0.02 (0.8523) | 1.97 (0.0826) | 0.02 (0.8523) |
| Homoerio-dictyol | 13.43 (<0.0001) | 66.29 (<0.0001) | 0.63 (0.0239) | 17.26 (<0.0001) | 0.03 (0.5882) | 0.71 (0.0177) | 0.02 (0.6937) |

### References

[1.] Gierczyk, B., Kaźmierczak, M., Popenda, Ł., Sporzyński, A., Schroeder, G., and Jurga, S. (2014) Influence of fluorine substituents on the NMR properties of phenylboronic acids, *Magn. Reson. Chem. 52*, 202-213. https://doi.org/10.1002/mrc.4051.

[2.] Lee, T. S., Krupa, R. A., Zhang, F., Hajimorad, M., Holtz, W. J., Prasad, N., Lee, S. K., and Keasling, J. D. (2011) BglBrick vectors and datasheets: a synthetic biology platform for gene expression, *J. Biol. Eng. 5*, 1-14. https://doi.org/10.1186/1754-1611-5-12.

[3.] Azubuike, C. C., Gatehouse, A. M., and Howard, T. P. (2021) pCAT vectors overcome inefficient electroporation of *Cupriavidus necator* H16, *New Biotechnol. 65*, 20-30. https://doi.org/10.1016/j.nbt.2021.07.003.

[4.] Dunstan, M. S., Robinson, C. J., Jervis, A. J., Yan, C., Carbonell, P., Hollywood, K. A., Currin, A., Swainston, N., Feuvre, R. L., Micklefield, J., Faulon, J.-L., Breitling, R., Turner, N. J., Takano, E., and Scrutton, N. S. (2020) Engineering *Escherichia coli* towards de novo production of gatekeeper (2*S*)-flavanones: naringenin, pinocembrin, eriodictyol and homoeriodictyol, *Synth. Biol. 5*, ysaa012. https://doi.org/10.1093/synbio/ysaa012.
